# The small cell lung cancer neuroendocrine transdifferentiation explorer

**DOI:** 10.1101/2022.08.01.502252

**Authors:** Ling Cai, Varun Sondhi, Mingrui Zhu, Esra Akbay, Ralph J. DeBerardinis, Yang Xie, John D. Minna, Guanghua Xiao, Adi Gazdar

## Abstract

SCLC is a high-grade neuroendocrine (NE) cancer that exhibits cellular plasticity. Transdifferention into non-NE cells creates considerable intra-tumoral heterogeneity, enhanced metastasis, greater tumor burden, and treatment resistance. Similar NE transdifferentiation has been observed in neuroblastoma (NBL). Targeting NE plasticity and cooperation between NE and non-NE cells in the tumor microenvironment may provide an avenue to enhance and restore sensitivity to available treatments. Although substantial transcriptomic changes take place upon NE transdifferentiation, conservation of these changes has not been investigated. In this study, we extensively curated genes associated with NE transdifferentiation in SCLC. We collected 35 datasets and compared the NE score-associated transcriptome across studies, for SCLC vs. NBL human tumors, human NBL tumors vs. cell lines, SCLC human tumors vs. tumors from genetically engineered mouse models (GEMMs), and SCLC GEMM uncultured cancer cells vs. cultured cancer cells. We have also created a user-friendly web application for researchers to explore these results. This work establishes a useful resource for researchers to understand the NE transdifferentiation landscape and explore context-dependent NE associations in SCLC and NBL.

## Introduction

Small cell lung cancer (SCLC) is a high-grade neuroendocrine (NE) malignancy with a dismal prognosis. 2 out of 3 patients with SCLC have extensive-stage disease at initial diagnosis, and over half of these patients die within a year. SCLC exhibits considerable inter- and intra-tumoral heterogeneity. Variant SCLC with loss of NE markers and increased expression of mesenchymal markers has been observed in 10-20% of SCLC cases and is associated with resistance to standard therapy [1]. Evidence from genetically engineered mouse models (GEMMs) suggests variant SCLC cells arise from transdifferentiation of NE to non-NE cells [2, 3], mimicking the NE deprogramming, expansion, and reprogramming to other cell types noted in a subset of pulmonary neuroendocrine cells in the setting of lung injury [4, 5]. The presence of non-NE malignant cells in the milieu enhances proliferation, metastasis, and chemoresistance of the malignant NE population [2, 6], suggesting the ability of NE cancer to produce stroma-like non-NE cancerous cells creates a self-sufficient favorable tumor microenvironment (TME) and negates the dependence on the non-cancerous stroma.

The stratification of malignant cells within SCLC tumors into NE and non-NE cells and the increasing appreciation of the plasticity between these subtypes has led to increased interest in targeting this plasticity and the cooperation between NE and non-NE cells [7]. Activation of the YAP/Notch/REST network [5] has been implicated in NE to non-NE transdifferentiation in native pulmonary neuroendocrine cells (PNECs) in normal lung tissue (reference), SCLC [2, 8, 9], neuroblastoma (NBL) [10-12], and pan-cancer studies [13, 14]. Substantial transcriptomic reprogramming takes place during this transition. Analysis of SCLC GEMMs has revealed significant changes in over one-fifth of the transcriptome between NE and non-NE cancer cells [5]. At the pathway level, loss of NE markers and activation of genes involved in cell adhesion, extracellular matrix, angiogenesis, and inflammation are the most prominent changes [5]. However, the full complement of pathway changes has not been enumerated in previous studies, and the conservation of the transdifferentiation program has not been assessed across different cancer types, between preclinical models and patient tumors, or between in vitro culture and the in vivo environment. In this study, we collected 35 datasets from SCLC, NBL, and normal lungs. Samples from these studies include bulk tumors from patients or GEMMs, patient-derived xenografts (PDXs), sorted tumor cells from GEMMs, and cultured cancer cells from patients or GEMMs. We explored and summarized the similarities and differences of NE and non-NE associated gene expression under different biological contexts. We also constructed a web-based application for users to review the results and perform their own ad hoc analyses.

## Results

### Transdifferentiation from NE to non-NE lineage involves substantial transcriptomic reprogramming

We evaluated the transcriptional association with the NE lineage by correlating gene expression values with NE scores computed from a 50-gene NE signature [8, 15]. We first evaluated the association in human primary SCLC tumors. Utilizing data from two large-scale human SCLC tumor studies, George_2015 [9] and Jiang_2016 [16], we performed meta-analyses to summarize the association between gene expression and NE scores. The transcriptome is evenly split between positively and negatively correlated genes (**Figure 1a**). We observed enrichment of negative correlation for 3,576 genes involved in innate immunity as denoted by innateDB[17] (**Figure 1b**), consistent with our previous findings that transition to the non-NE lineage abrogates repression of immune gene expression [15]. In addition, we performed an extensive manual review of the NE score-correlated gene results. Out of the 15,792 genes analyzed by meta-analyses, 3,771 (24%) genes exhibited a medium or strong association (absolute r > 0.3) with NE score, and 935 of these genes were curated into 47 families/pathways (**Figure 1c**). Among these, 10 families/pathways were further classified into 160 subgroups. **Figure 1d** provides an example, showing the NE association profiles for eleven subgroups under “Metabolic pathways”. We provide visualizations of subgroup NE association profiles from the other nine families/pathways in **Figure S1**. In our curated lists, the majority (1106, 94%) of genes are significantly correlated genes (absolute r > 0.165, padj < 0.05), but at times we have also included other genes in the same pathway and genes known to be important for SCLC biology. The full lists of curated pathways are provided in **supplementary information 1** in alphabetical order. **Figure 1e** provides a snapshot of one of the subgroup tables. We also dichotomized samples from tumor or SCLC cell line datasets into NE and non-NE groups and calculated the standardized mean difference between the two groups. Furthermore, from a high-NE score (0.91) H69 cell line, we derived a low-NE score (−0.02) adherent H69-AD cell line and compared RNA-seq data from both lines. We verified the loss of selected NE markers (Ep-CAM, NCAM, and DLL3), and gain of selected non-NE markers (CD44 and HLA) by flow cytometry and demonstrated good concordance with the RNA-seq data (**Figure S2**). The FPKM difference was calculated and compared to the FPKM difference for NE and non-NE cell line groups. These data are incorporated into each summary table and colored for easier visualization; to examine the consistency between tumor data, cell line data, and our paired cell line model (**Figure 1e** and **supplementary information 1**).

**Figure 1.**
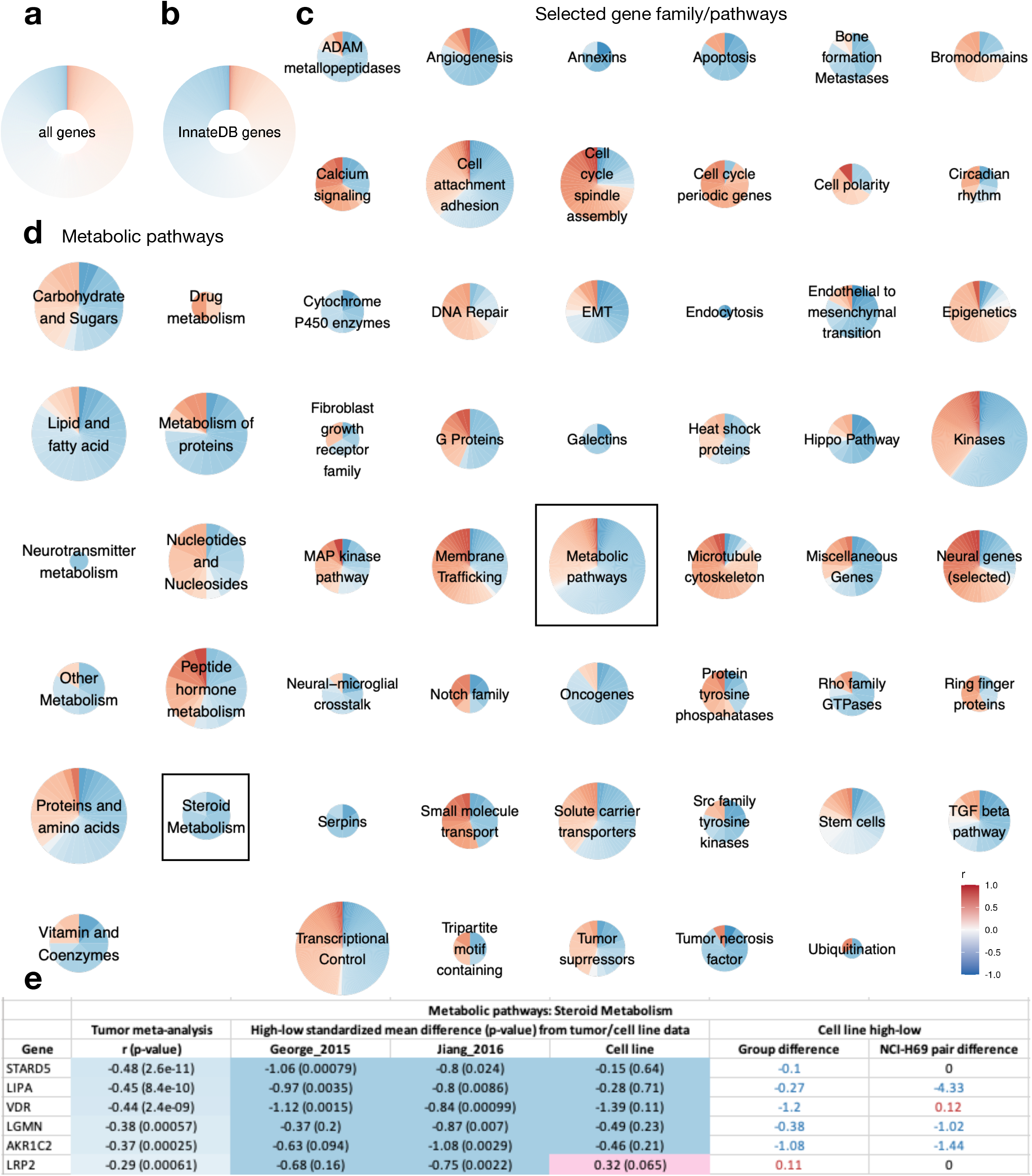
Dissecting pathways associated with NE-to-non-NE transdifferentation. **a**. Correlation between NE scores and gene expression across 15,792 genes in SCLC tumor datasets. These genes are evenly split between positively and negatively correlated genes. **b**. Correlation between NE scores and expression of 3,576 innate immune genes from InnateDB. Over 60% of the genes are negatively correlated with NE scores. Among the genes with a large effect size (absolute r > 0.5), 80% are negatively correlated with NE score. **c**. NE score correlation in 47 selected gene families/pathways related to SCLC biology. **d**. NE score correlation for 11 subsets of genes under “Metabolic pathways”. For **c** and **d**, pie sizes are proportional to the number of genes within the set. **e**. Snapshot of table summarizing NE score correlation for gene members of “Steroid Metabolism” (see also **supplementary information 1**). George_2015 and Jiang_2016 are two SCLC tumor studies. K-means clustering was used to determine NE and non-NE groups for the calculation of standardized mean difference. In the “Cell line high-low” section of the table, FPKM difference between NE and non-NE groups, or H69 parental and H69 adherent derived cell line was reported.

From our manual review, we identified many NE score-associated genes from pathways well known to be important for neural and endocrine functions, such as neural genes, microtubule cytoskeleton, and membrane trafficking genes. For genes negatively associated with NE scores, we found genes known to be important for mesenchymal functions, such as epithelial-to-mesenchymal transition (EMT), endothelial-to-mesenchymal transition, angiogenesis, bone formation, metastasis, and stem cell function. Many cell cycle genes positively correlate with NE scores, whereas many apoptosis genes and tumor necrosis factors negatively correlate with NE scores. Several gene families have multiple members that exhibit concerted anticorrelation with NE scores, such as proteases from the ADAM metallopeptidases and serpins families, cytochrome P450 enzymes, the lipid-binding protein annexins, and the carbohydrate-binding protein galectins (**Figure 1c**). Our subgroup classification provides a more in-depth examination of large families/pathways such as transcriptional control and kinases (**Figures S1a**-**b**). Sometimes, most of the subgroups followed the same trend as the parental group. For example, most of the lysine demethylase, lysine methyltransferase, polycomb complex, and SWI/SNF complex genes under “Epigenetics”, and most of the kinesin, stathmin, and tubulin genes under “Microtubule cytoskeleton” exhibit positive correlations with NE scores (**Figures S1 c-d**). Occasionally opposite trends are observed for different subgroups of the parental group. For example, under “Small molecule transport”, potassium channel and sodium channel genes positively correlate with NE scores, whereas aquaporins and S100 proteins negatively correlate with NE scores. Under “Cell attachment adhesion”, the majority of the protocadherins, desmosomes, and synapse genes positively correlate with NE scores, whereas integrins, lectins, and fibulins negatively correlate with NE scores (**Figures S1e** and **h**)

### Comparison of NE to non-NE lineage transcriptomic changes across different cancer types, sample types, species, and culture conditions

Having noted substantial transcriptomic changes associated with NE to non-NE transition in SCLC tumors, and some disagreement between SCLC tumors and SCLC cell lines (**supplementary information 1**), we wanted to further explore NE and non-NE associated gene expression under different biological contexts. We expanded our data collection to include additional SCLC human tumor, SCLC PDX, and GEMM datasets, as well as NBL human cancer cell line and tumor datasets. We computed the NE score correlation coefficient for the whole transcriptome in each dataset and hierarchically clustered the 35 datasets based on these coefficients (**Figure 2**). Interestingly, while positive correlations were observed for almost all dataset pairs, indicating considerable conservation, the unsupervised clustering approach also spontaneously organized the datasets into five clades with distinct properties – one for human SCLC samples (including cell line, PDX, and patient tumors), one for uncultured SCLC GEMM samples, one for cultured SCLC GEMM cell lines, one for NBL human cell lines, and one for NBL human tumors (**Figure 2**). We next performed a series of four comparisons: human SCLC vs. NBL tumors, human NBL cell lines vs. tumors, human SCLC vs. GEMM tumors, and SCLC GEMM sorted vs. cultured cancer cells. These comparisons allowed us to understand the context-specific NE score associations in more detail. The comparison scheme is illustrated in **Figure S3a**. For each comparison, we performed meta-analyses for datasets from each side and compared the summary correlation. We identified set-specific genes with robust correlation in one set (absolute r > 0.4), but weak (absolute r < 0.1) correlation in the opposite direction in the other set. We also identified genes concertedly correlated or anticorrelated with NE scores in both sets (**supplementary information 2-5, sheet 1** and **Figures S3b-e**). We then performed pathway enrichment analyses with eleven gene family/pathway libraries for these inconsistent or consistent genes to understand the underlying biology (**supplementary information 2-5, sheet 2-5**).

**Figure 2.**
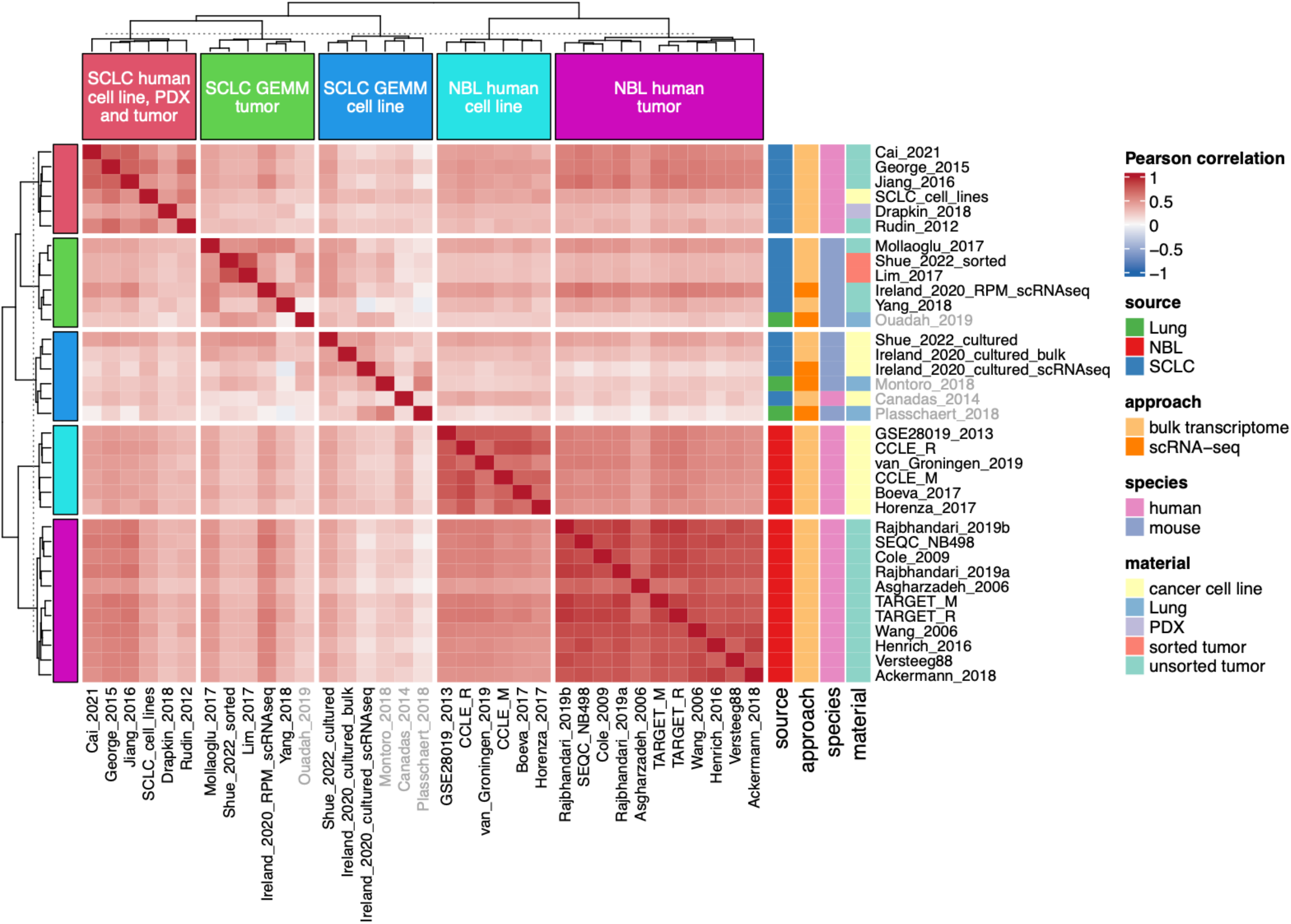
Transcriptomic changes from NE to non-NE lineage transdifferentiation are conserved between SCLC and NBL, in preclinical models and patient tumors. Hierarchically clustered correlation heatmap that spontaneously organizes datasets by source, species, and material. For each dataset, the correlations between NE score and gene expression across the whole transcriptome were computed. Pairwise correlations for study pairs were then calculated to generate this correlation heatmap. Positive correlations (red) were observed for almost all of the study pairs, suggesting conservation of NE transcriptome under these different contexts. Although the datasets were generated by many different platforms, studies with the same source, species, and material cluster tighter together, suggesting distinctive NE-associated features also exist under different contexts. In the heatmap, studies with properties different from the clade label are marked in gray.

#### A comparison between SCLC and NBL in human patient tumors

Although SCLC originates from resident neuroendocrine cells in the lung, and NBL originates from neural crest cells commonly in and around the adrenal glands, they are both NE malignancies and have both been observed to transition to a non-NE fate [18, 19]. The NE and non-NE genes from the existing SCLC-based 50-gene NE signature exhibited robust differential expression in the NBL cell lines and tumors. (**Figures S4a**-**b**). However, the expression of certain genes is low or missing in some of the NBL datasets. We hence derived an NBL-specific 50-gene NE signature (**supplementary information 6**) to compute NE scores from NBL datasets (**Figures S4c**-**d**). We compared the NE score-associated transcriptome in four human SCLC tumor studies and eleven human NBL tumor studies (**Figure S5**). We performed pathway enrichment analyses (**Figures S5a**-**c**), examined consistency across studies for significantly changed genes from top hits (**Figures S5d**-**e**), and examined patterns of gene expression from selected studies (**Figure S5f**). Among genes with consistent NE score associations in both SCLC and NBL tumor datasets, positively correlated genes are enriched for neural genes and negatively correlated genes are enriched for immune genes. On the other hand, the SCLC-specific or NBL-specific NE score-associated genes are enriched in organ development or cell differentiation (**Figures S5c** and **S5e**). Many of these lineage-specific genes are transcription factors (TFs), including *ASCL1, NKX2-1*, and *NKX2-2*, which are known to be important for SCLC; and *HAND2, GATA3*, and *PHOX2B*, which are known to be important for NBL (**Figure 3a**). Other TFs including the sine class homeoboxes, NKL subclass homeoboxes, Forkhead boxes, and basic helix-loop-helix proteins (**Figure S5a**,**d**, and **Figure 3b,c**) were also found to be uniquely associated with NE scores in SCLC or NBL. We also found cancer-type-specific NE-associated metabolic gene expression. Several fatty acid metabolism genes are only upregulated in low-NE score SCLC tumors, whereas several glutathione S-transferases are only upregulated in low-NE score NBL tumors (**Figures S5a**,**b**, and **3b**,**c**). Among the cell cycle genes, down-regulation of *CCND1* and *CDK6* and upregulation of *CDK2* and *CDKN2A* in high-NE score tumors are also unique in SCLC, likely due to the frequent loss of *RB1* in SCLC but not NBL (**Figure 3b**). We noted that low-NE score samples are rare in NBL tumor sets but frequent in NBL cell line sets (**Figure S4**). We hence sought to identify genes that consistently associate with NE scores in SCLC tumors and NBL cell lines but not NBL tumors. We found 49 genes that met such criteria (**Figure 3d**), including NE driver TF *ASCL1* and non-NE marker *CD44*. These results suggest NE transdifferentiation is more prominent in cell lines than in patient tumors.

**Figure 3.**
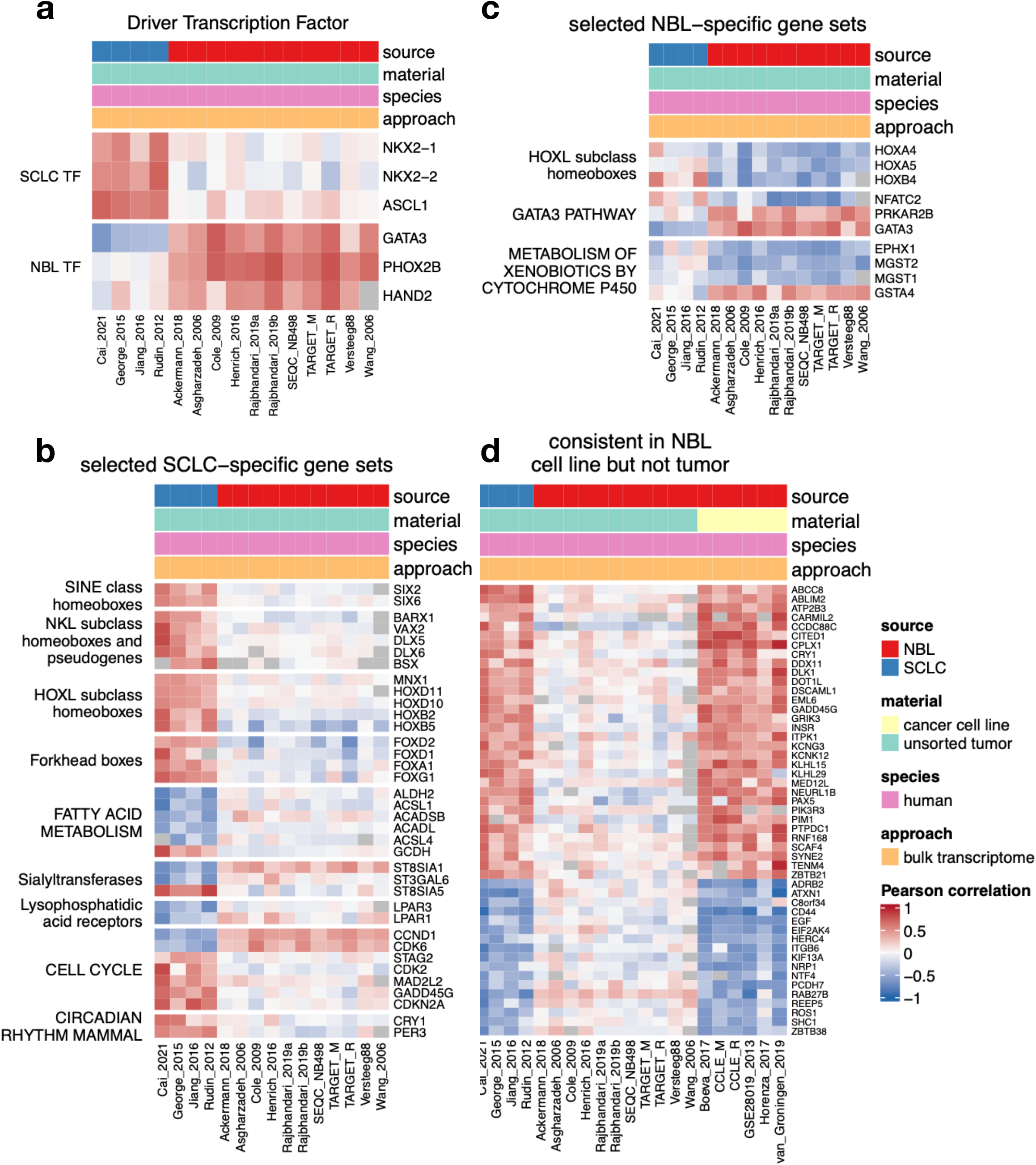
Selected SCLC- or NBL- specific NE score - gene expression correlation heatmaps. **a**. Distinct NE score associations for driver TFs in SCLC and NBL. **b**. Selected SCLC-specific NE score associations. **c**. Selected NBL-specific NE score associations. **d**. NE score associations consistent between SCLC tumors and NBL cell lines but not NBL tumors.

#### A comparison between human NBL cell lines and tumors

We compared the NE score-associated transcriptome in six human NBL cell line studies and eleven human NBL tumor studies (**Figure S6**). We again noted consistent enrichment in neural and immune defense genes for both tumor and cell line datasets, suggesting cell-autonomous NE regulation. But we also noticed many tumor-specific anticorrelation between immune genes and NE scores, suggesting the presence of non-cell-autonomous immune gene regulation (**Figures S6c**,**e**). Many innate and adaptive immune genes were found to negatively associate with NE scores in tumors but not cell lines (**Figure 4a**). When we compared the NE score association for genes curated by innateDB in tumors vs. cell lines, we observed a similar pattern in NBL and SCLC – while many genes distribute along the diagonal line, a large number of genes have robust negative correlations with NE scores only in tumors but not in cell lines (**Figure 4b**). For genes specifically associated with NE scores in cell lines, we found many N-glycan biosynthesis genes (**Figure S6b**), glycoproteins PGS4, PGS5, and PGS9 (**Figure S6d**,**f**) have higher expression in low-NE score NBL cell lines. Several genes involved in glycosylphosphatidylinositol biosynthesis, phosphatidylinositol signaling, and clathrin adaptor complex subunits also exhibited differential association with NE scores in cell lines but not tumors (**Figure 4c**).

**Figure 4.**
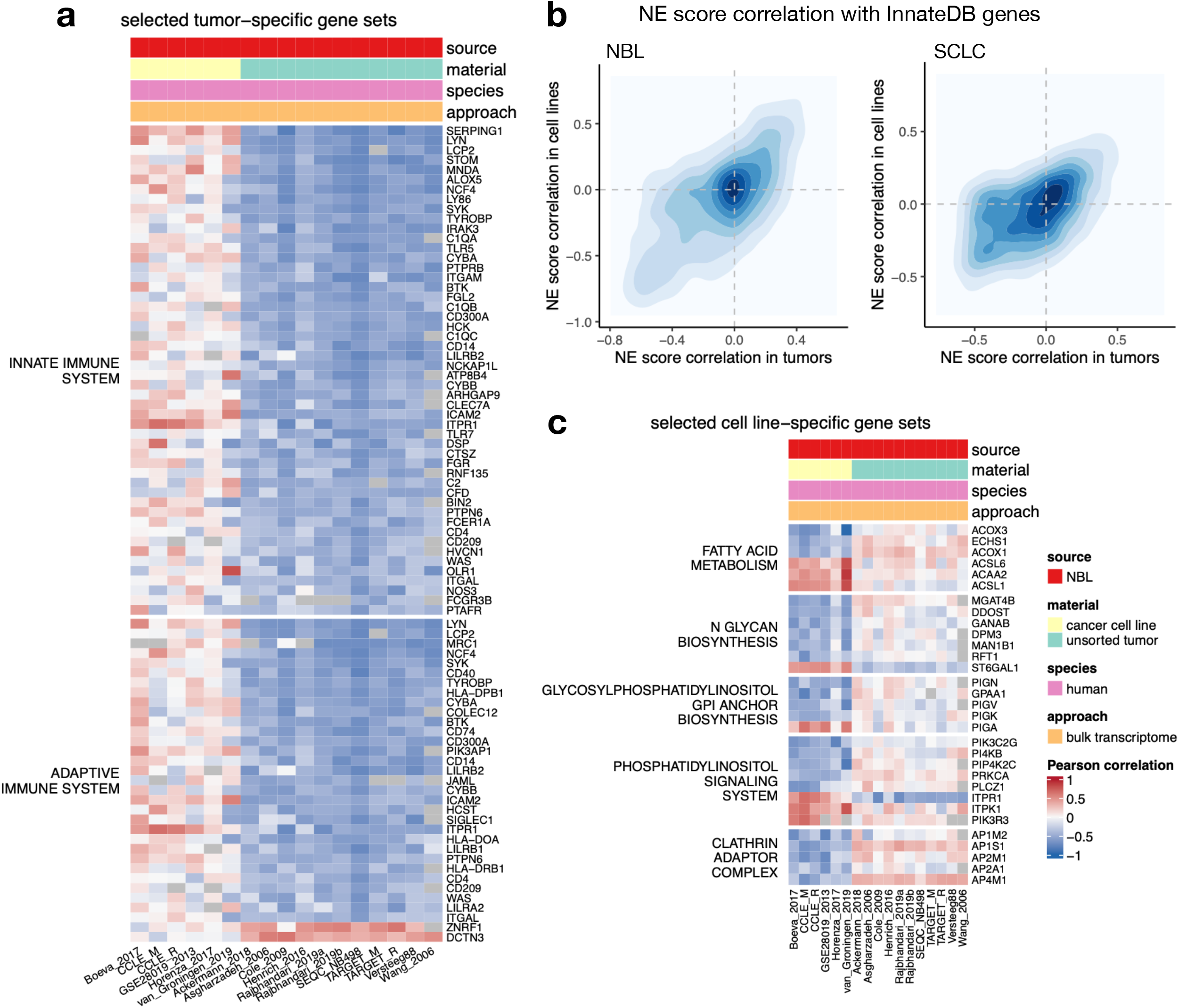
Selected NBL cell line- or tumor- specific NE score - gene expression correlations. **a**. Genes from the innate and adaptive immune system genesets that correlate with NE scores in tumors but not cell lines. **b**. Comparing tumor (x-axis) vs. cell line (y-axis) NE score correlation in NBL (left) and SCLC (right) for innate immune genes curated by InnateDB. Correlation coefficients are from meta-analyses in NBL, as specified in Figure S2a. SCLC tumor NE score correlation comes from meta-analysis of George_2015 and Jiang_2016 datasets, whereas SCLC cell line NE score correlation comes from a single dataset. **c**. Selected NBL cell line-specific NE - gene correlations.

#### A comparison between human SCLC tumors and GEMM tumors

We compared the NE score-associated transcriptome in four human SCLC tumor studies and two SCLC GEMM tumor studies (**Figure S7**). While genes negatively associated with NE scores in both SCLC human and GEMM tumors continue to be most significantly enriched for immune defense; cell cycle genes rather than neural genes have the most significant enrichment with positive NE score association from both human and GEMM SCLC tumors (**Figure S7**). We found that for some of the human-specific NE score-associated genes, such as the granins, NE score association in the GEMM dataset “Mollaoglu_2017” was more consistent with the human data than in “Yang_2018” (**Figure 5a**). The 45 tumors from “Yang_2018” are all from Rb/p53/p130 (TKO) mice and all have a very high NE score, whereas two of the Rb/p53/Myc (RPM) tumors from “Mollaoglu_2017” have negative NE scores (**Figure 5b**). The NE score associations with neuronal development genes are more positive in “Mollaoglu_2017”, as in the human data (**Figure 5c**). Human tumors exhibit significant inter-tumoral heterogeneity; our analysis suggests the lack of inter-tumoral heterogeneity in GEMM datasets can confound a comparison of NE score associated genes in human specific models with NE score associated genes in the GEMMs. On the other hand, consistent correlations between mouse-specific NE score-associated genes are noted across GEMM datasets, largely outnumbering the genes noted in the other three categories – human tumor specific genes, genes that are positively correlated in human tumors and GEMM tumors, and genes that are negatively correlated in human tumors and GEMM tumors (**Figure S3d**). Many of these mouse-specific genes are involved in metabolism, such as sphingolipid metabolism, fatty acid catabolism (negative correlations), and purine nucleoside biosynthesis (positive correlations) (**Figure 5d**). Many cell-cell junction proteins and keratin-related proteins also exhibit concerted negative correlations with NE scores in GEMM tumors (**Figure 5d**). These include many cadherin and claudin genes, suggesting in GEMM tumors the low-NE score samples are more epithelial-like.

**Figure 5.**
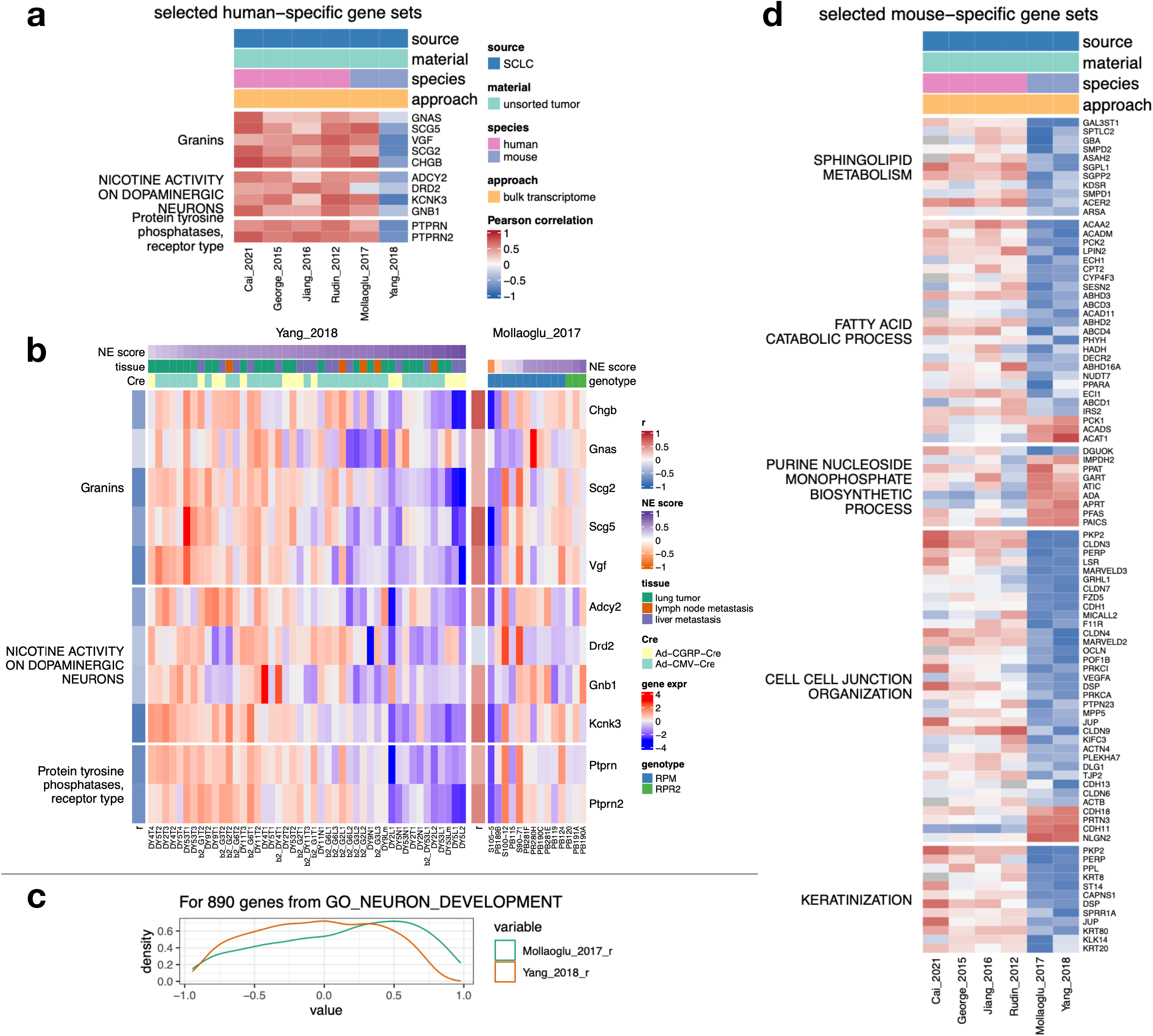
Selected SCLC human- or mouse- specific NE score - gene expression correlations. **a**. For some human tumor-specific NE score associated genes, GEMM dataset Mollaoglu_2017 produced more agreeable correlations than Yang_2018. **b**. Yang_2018 GEMM samples have lower inter-tumoral heterogeneity for NE scores than samples from Mollaoglu_2017. **c**. Mollaoglu_2017 dataset exhibits more positive NE score correlations for genes related to neuron development. **d**. Selected metabolic and structural genes that exhibit mouse-specific NE score correlations.

#### A comparison between SCLC GEMM uncultured and cultured cancer cells

We compared the NE score-associated transcriptome in two SCLC GEMM “uncultured” datasets and two SCLC GEMM “cultured” datasets (**Figure S8**). In these studies, TKO NE and non-NE SCLC cells were isolated by flow cytometry and profiled directly or further cultured as cell lines[2, 5], whereas RPM SCLC cells were isolated and grown in vitro. NE-to-non-NE transition is induced as RPM cells adapt to in vitro culture [3]. As there is significant NE heterogeneity among these samples, larger effect sizes in NE association could be observed with these datasets than in the bulk tumor/cell line datasets (**Figure S9a**). Of note, both uncultured datasets were generated from the same TKO GEMM by the same research group[2, 5] whereas the two cultured datasets were generated from different GEMMs (TKO vs. RPM) by different groups[3, 5]. We observed better concordance for the uncultured datasets compared to the cultured datasets (**Figure S9b**-**c**). Nevertheless, the datasets still cluster by culture status based on NE score-transcriptome correlations (**Figure S9d**). Many NE score-gene expression associations appear to be specific for uncultured cancer cells (**Figure S2e**). Strikingly, close to one-third of the solute carrier (SLC) gene family members (105/326) appear to have different NE score associations in uncultured and cultured conditions (**Figures S8a** and **6a**). From the pathway analysis result using gene ontology biological process library (GO BP), we also identified 25 gene sets related to “transport” (**supplementary information 5**). Of the 306 GO BP annotated genes that were found to be uncultured-specific, 90 are SLC genes. Of the 149 genes with better consistency between the two cultured datasets, 49 are SLC genes (**Figures S9e**). These results suggest metabolite transport is differentially regulated in NE and non-NE cells in vivo, but many of these differences are lost when cells are cultured in vitro. We also found robust negative NE score correlations for interferon-gamma signaling genes and surfactant genes in uncultured sorted SCLC cells, but not in cultured cells, suggesting the expression of these genes in non-NE cancer cells is dependent on factors in the TME and is not recapitulated in vitro. Similarly, many glycolytic genes and some glutamate/glutamine metabolism genes are positively correlated with NE scores in sorted cells but not cultured cells, suggesting differential carbon and nitrogen regulation in NE and non-NE cells in vivo, which is lost upon culturing these cells in vitro (**Figure 6b**). Among culture-specific NE score-associated genes, we found many involved in epithelial cell differentiation (**Figure 6c**). Many apical junction assembly genes appear to have distinct NE score associations in uncultured vs. cultured conditions (**Figure 6b**). We specifically examined several genes related to EMT or from the Myc gene family. We evaluated these genes in SCLC GEMM uncultured and cultured cells from Shue_2022[5] as well as human SCLC cell line H69 and H69M from Canadas_2014[20] (**Figure 6c**). Among the mesenchymal markers, *CD44/Cd44, FN1/Fn1* are consistently upregulated in low-NE score uncultured and cultured samples, while *ZEB1/Zeb1, VIM/Vim*, and ANXA1/*Anxa1* only exhibited changes in cultured samples (**Figure 6d**, top). Higher expression of epithelial markers *CLDN8/Cldn8* and *CLDN10/Cldn10* in non-NE cells can only be observed in uncultured SCLC GEMM cells but not in the cultured samples (**Figure 6d**). For Myc family members, we observed increased Myc expression in low-NE score samples of cultured SCLC cells from both human tumors and GEMMs, but not uncultured SCLC GEMM cells. Higher Mycl levels were found in cultured high-NE score SCLC GEMM cells, but not in human H69 or uncultured SCLC GEMM cells (**Figure 6d**, bottom). These results suggest the culture condition of cells can have a profound impact on the expression of EMT genes and Myc genes in non-NE SCLC cells.

**Figure 6.**
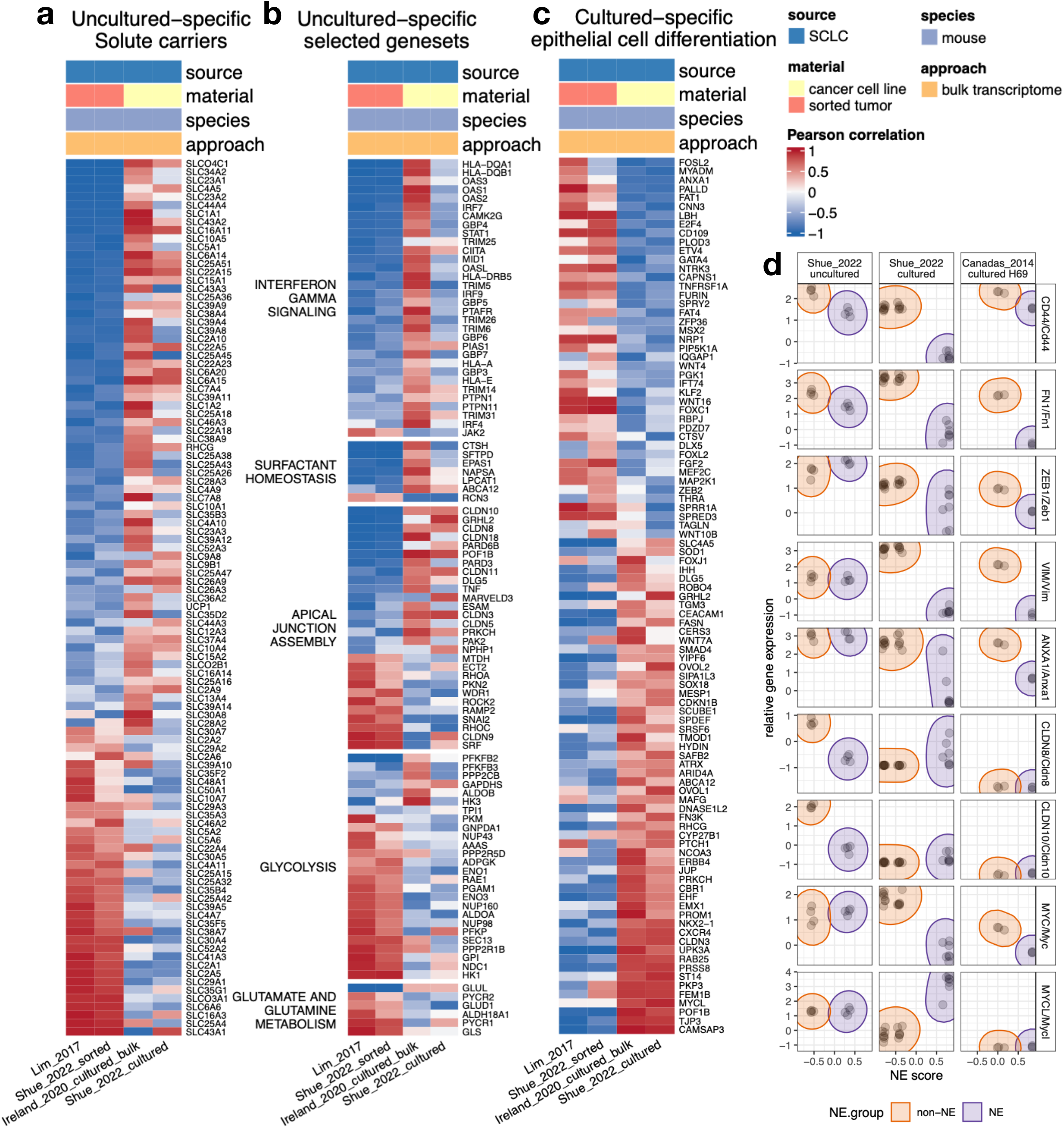
Investigation of gene - NE score associations specific to culture conditions with SCLC GEMM datasets. **a**-**b**. Genes with strong NE score associations specifically under the uncultured condition, including members of the Solute carrier (SLC) family (**a**) and other selected pathways (**b**). **c**. Epithelial cell differentiation genes with strong NE score associations specifically under the cultured condition. **d**. Comparison of NE association for selected mesenchymal/epithelial markers and Myc family genes in human NE and non-NE SCLC cell lines, SCLC GEMM NE and non-NE cell lines as well as uncultured SCLC GEMM NE and non-NE cells.

### An SCLC NE transdifferentiation explorer

Beyond the results included in this paper, with the understanding that researchers may want to perform their own analyses to explore NE score-associated transcriptional changes, we constructed a web application at https://lccl.shinyapps.io/GSNE/ (**Figure 7**). Detailed tutorials and instructions are provided in the web application on how to use the tools. In the *Cross-study summary* tool, users may review the correlation between NE score and gene expression (**Figure 7a**). This is provided for 35 individual studies, but we also performed six sets of meta-analyses to summarize correlations from datasets with similar properties. The datasets included in each set of meta-analyses essentially follows the scheme we used in **Figure S3a**, except that we additionally included one SCLC PDX dataset and one SCLC human cell line dataset for the meta-analyses “SCLC human cell lines, PDX, and tumors”. When using this tool, users may select different results to create a big table joined by genes. Users may review the full table or genes from pre-defined gene sets (**Figure 7a**). In the *Pathway analyses* tool, users may perform hypergeometric tests for top genes positively or negatively correlated with NE scores using nine gene set libraries (**Figure 7b**). In the *Heatmap* tool, users may select from 30 datasets to construct heatmaps. The datasets can be selected by source (NBL or SCLC), species (human or mouse), and sample type (unsorted tumor, sorted tumor, cancer cell line, or PDX). Users may use pre-defined genes from our curation or published gene set libraries. We have additionally included a few signature gene sets, including our SCLC and NBL NE gene signature, as well as an interferon gamma signature that predicts response to immunotherapy[21]. Or users may enter genes of their interest. Note that in the *Pathway Analyses* tool, the “Overlapping Genes” were provided as comma-delimited lines. Users may copy and paste these gene lists to visualize them in a specific dataset using the Heatmap tool. Sample annotation with NE score and SCLC TF expression status can be added to the heatmap and be used to order the samples. By default, expression values are z-transformed for each gene, but users may also generate heatmaps with untransformed data. Lastly, the annotation and sources of these datasets are listed in the *Reference* tool (**Figure 7d**).

**Figure 7.**
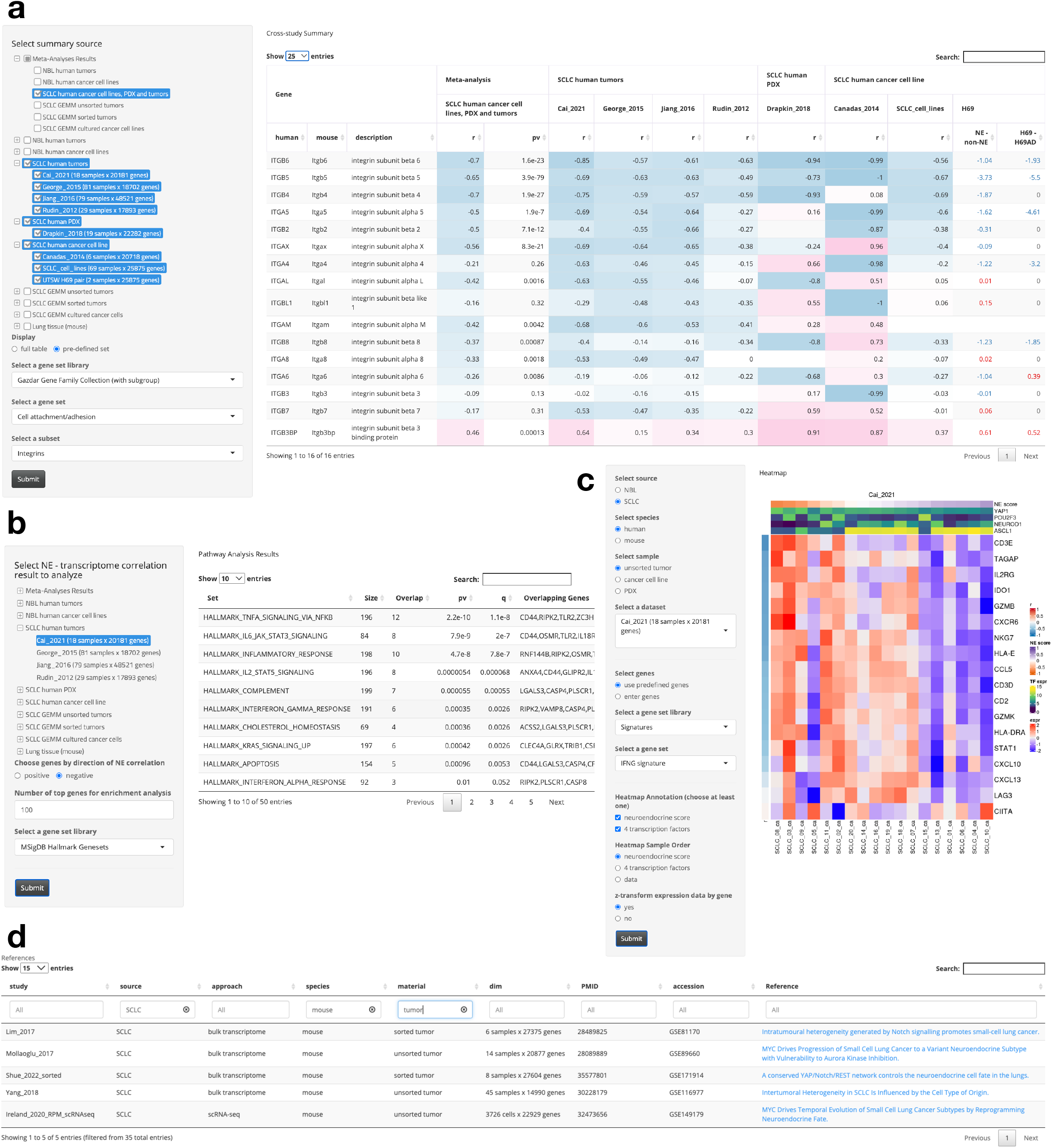
A web application for exploring NE - non-NE transdifferentiation-associated gene expression changes. **a**. Cross-study Summary tool. Users may review correlation between gene expression and NE scores from individual studies or meta-analysis summarized results as an outer-joined table. Users may review the full table or a set of pre-defined genes. **b**. Pathway Analyses tool. Users may select NE score - transcriptome correlation results from an individual study or a set of meta-analysis to perform hypergeometric tests for top positively or negatively correlated genes. **c**. Heatmap tool. Users may choose from a total of 31 datasets by specifying source, species and sample material. A pre-defined geneset or user specified geneset can be used for heatmap generation. In the heatmap, samples can be further annotated with NE scores and expression of SCLC driver TFs; genes are annotated by their correlation with NE score. **d**. Reference interface. The reference list all 36 studies used for this web application. Studies can be classified by source, approach and material. Sample sizes and number of genes profiled for each study are provided. Data accession ids and original publications are also provided. Tutorials are also available on https://lccl.shinyapps.io/GSNE/.

## Discussion

We collected and compared 35 datasets from various biological sources including SCLC patient derived tumors, SCLC PDXs, SCLC cell lines, SCLC sorted GEMMs, SCLC cultured GEMMs, NBL tumors, and NBL cell lines. While overall conservation of NE score associations could be observed across datasets from different cancer types, species, and sample types, we also found consistent context-specific NE score associations in sample sets with similar biological properties (**Figure 2**). In the comparison between SCLC and NBL, we identified distinct sets of TFs that associate with NE scores (**Figures 3a-c**), implying distinct interplays between NE TFs and other lineage TFs in SCLC and NBL. Unlike the SCLC tumor datasets, low-NE score samples are rare in NBL tumor datasets (0-3% negative NE score samples). In contrast, low-NE score samples are more frequent in NBL cell lines (5-24%) (**Figure S4**). The underlying cause of this difference remains unclear, and one can hypothesize that non-NE NBL lines could be easier to establish in culture, or that in vitro culture conditions increase the differentiation from a NE to a non-NE state, as has been shown in the RPM SCLC GEMM[3]. In the comparison between cell lines and tumors, we identified many tumor-specific immune gene correlations in NBL as well as SCLC (**Figure 4b**), suggesting significant immune modulation by the TME.

As the NE transdifferentiation process in SCLC resembles the NE transdifferentiation of PNECs in normal lung, we were also interested in the NE score associated transcriptome from the normal lung and included three mouse scRNA-seq datasets: “Montoro_2018”, “Plasschaert_2018”, and “Ouadah_2019” [4, 22, 23]. However, due to the high dropout rate of scRNA-seq data, the correlations were weak in these datasets. We therefore did not perform an in-depth analysis of these datasets but have provided the NE score correlation results in the web application.

Past studies of SCLC GEMMs have provided valuable insights into intra-tumoral heterogeneity of SCLC. We observed interesting metabolic NE score associations in SCLC GEMMs. However, many associations observed in GEMM tumors were not seen in human tumors, such as those for genes from the purine nucleoside biosynthesis and fatty acid metabolism pathways (**Figures 5d**). Some metabolic associations were only observed in uncultured GEMM SCLC cells but not in cultured GEMM SCLC cells, these include many of the metabolite transporters, glycolytic genes, and nitrogen catabolism genes (**Figures 6a-b**). These unique findings from uncultured GEMMs not observed in cultured GEMM cells suggests some SCLC metabolic properties are unique to the in vivo setting and are lost when tumor cells are cultured in vitro. A similar effect remains to be validated in human studies due to the paucity of data that would allow a direct comparison of the transcriptome of primary human tumors to cultured counterparts. Although the human tumor atlas network (HTAN) consortium has published human SCLC scRNA-seq data for 21 patient specimens [24], the HTAN cohort did not include YAP1+ low-NE score tumors and the only POU2F3+ tumor (RU1322) has predominantly high-NE score malignant cells. We found two of the NEUROD1+ tumors (RU1215 and RU1231) exhibited some degree of intra-tumoral heterogeneity, but the low-NE score cells are rare in the malignant cell population of these samples and the resulting NE score associations are weak. We thus excluded the results from HTAN in our collection.

In summary, in this work we curated an extensive collection of NE score-correlated genes in SCLC and examined the conservation and differences of NE transdifferentiation programs in different systems. We constructed a web application for researchers to explore NE score-associated gene expression in 35 datasets across a wide array of SCLC and NBL models. We hope this becomes a useful resource for researchers to explore and better understand the NE transdifferentiation landscape. This work and the associated web tool can aid researchers studying NE plasticity, intratumoral heterogeneity, and cooperation between NE and non-NE cells in the TME. It should also serve as a valuable resource for those working to devise new therapeutic strategies to target NE plasticity and intercellular cooperation in the TME.

## Material and methods

### NE score calculation

The original SCLC NE signature based on microarray gene expression data was described in Zhang et al. [8]. Here, we use the updated signature generated from RNA-seq expression[15]. We also generated an NE signature for NBL. We first generated NE scores based on the SCLC NE signature for six NBL cell line datasets. We then computed NE score – gene expression correlation for these datasets and used meta-analysis to generate summary correlation values. We examined genes that are covered in all six NBL cell line datasets and eleven NBL tumor datasets. The top 25 genes positively correlated with NE score and top 25 genes negatively correlated with NE scores are selected to generate the NBL NE signature. Instead of average expression values from a given dataset that was used in the SCLC NE signature, we assigned equal weights of 1 for each of the NE and non-NE signature genes in the NBL signature. A quantitative NE score can be generated from an NE signature using the formula: NE score = (correl NE – correl non-NE)/2, where correl NE (or non-NE) is the Pearson correlation between expression of the 50 genes in the test sample and expression/weight of these genes in the NE (or non-NE) cell line group. This score has a range of −1 to +1, where a positive score predicts for NE while a negative score predicts for non-NE cell types. The higher the score in absolute value, the better the prediction.

### Flow cytometry

Construction of the adherent H69-AD subline from the parental H69 cell line has previously been described[15]. Cell lines were first stained with fixative dead cell stain (Fisher Cat# 50-112-1528) RT for 8min to gate the live cells. Then they were stained with fluorophore conjugated antibody or isotype for 20min on ice. Samples were washed with FACS buffer (2% FBS in PBS) after every incubation with fixative dye or antibody. All antibodies were diluted in FACS buffer. All samples were analyzed by BD FACS Canto. Results were analyzed in FlowJo software.

| Antibody | Vendor | Cat. # | Isotype Cat. # |
| --- | --- | --- | --- |
| PE anti-human HLA-A,B,C Antibody | Biolegend | 311406 | 400213 |
| Pacific Blue™ anti-mouse/human CD44 | Biolegend | 103020 | 400627 |
| Human DLL3 PE-conjugated Antibody | Fisher: R&D | FAB4315P | IC108P |
| APC anti-human CD56 (NCAM) Antibody | Biolegend | 362504 | 400121 |
| PerCP/Cy5.5 anti-human CD326 (Ep-CAM) | Biolegend | 324214 | 400337 |

### Data processing

The sources of the datasets have been listed in the “Reference” section of https://lccl.shinyapps.io/GSNE/. Datasets from the Gene Expression Omnibus (GEO) were downloaded using R package GEOquery[25]. CCLE RNA-seq (CCLE_depMap_19Q1_TPM.csv) and microarray (CCLE_Expression_Entrez_2012-09-29.gct) datasets were downloaded from the DepMap portal[26]. For datasets from Therapeutically Applicable Research to Generate Effective Treatments (TARGET) (https://ocg.cancer.gov/programs/target) initiative, phs000467 [27], “TARGET_M” for microarray data was downloaded from the TARGET Data Matrix, whereas “TARGET_R” for RNA-seq data was downloaded from the UCSC Toil RNAseq Recompute Compendium [28]. George_2015 dataset comes from supplementary file from the original paper[9], “Rudin_2012” dataset is from the authors of the original paper[29]. Genesets used for analyses and we bapplication were downloaded from the Molecular Signature Database (MSigDB)[30]. All analyses were conducted in *R*[31].

### Development of web application “GSNE”

The web application https://lccl.shinyapps.io/GSNE/ is a shiny app deployed at the shinyapps.io servers. It is implemented through the following R packages: ‘data.table’, ‘DT’, ‘ComplexHeatmap’, ‘viridis’, ‘RColorBrewer’, ‘shiny’,’ slickR’, ‘shinyjs’, ‘shinythemes’, ‘shinyTree, and ‘shinycssloaders. The source code is be available at a GitHub repository https://github.com/cailing20/GSNE.

## Acknowledgment

Our web application “<u>G</u>azdar <u>S</u>CLC <u>N</u>E <u>E</u>xplorer (GSNE)” is named after Dr. Adi Gazdar in remembrance of his mentorship and contribution to SCLC research. L.C. receives support from UTSW ACS-IRG (IRG-21-142-16) and a Lung Cancer SPORE Career Enhancement Program award from P50CA70907. This study is supported by funding from the National Institutes of Health [R35CA22044901, P30CA142543, P50CA70907, and R35GM136375], and the Cancer Prevention and Research Institute of Texas [RP190107 and RP180805]. R.J.D. receives funding from Howard Hughes Medical Institute. This article is subject to HHMI’s Open Access to Publications policy. HHMI lab heads have previously granted a nonexclusive CC BY 4.0 license to the public and a sublicensable license to HHMI in their research articles. Pursuant to those licenses, the author-accepted manuscript of this article can be made freely available under a CC BY4.0 license immediately upon publication. R.J.D. is an advisor for Agios Pharmaceuticals and Vida Ventures and a co-founder of Atavistik Bio. JDM receives licensing fees from the NIH and UTSW for distribution of human tumor cell lines. The other authors declare no competing interests.

## Reference

1. Carney, D.N., J.B. Mitchell, and T.J. Kinsella, In vitro radiation and chemotherapy sensitivity of established cell lines of human small cell lung cancer and its large cell morphological variants. Cancer Res, 1983. 43(6): p. 2806–11.

2. Lim, J.S., et al., Intratumoural heterogeneity generated by Notch signalling promotes small-cell lung cancer. Nature, 2017. 545(7654): p. 360–364.

3. Ireland, A.S., et al., MYC Drives Temporal Evolution of Small Cell Lung Cancer Subtypes by Reprogramming Neuroendocrine Fate. Cancer Cell, 2020.

4. Ouadah, Y., et al., Rare Pulmonary Neuroendocrine Cells Are Stem Cells Regulated by Rb, p53, and Notch. Cell, 2019. 179(2): p. 403–416 e23.

5. Shue, Y.T., et al., A conserved YAP/Notch/REST network controls the neuroendocrine cell fate in the lungs. Nat Commun, 2022. 13(1): p. 2690.

6. Calbo, J., et al., A functional role for tumor cell heterogeneity in a mouse model of small cell lung cancer. Cancer Cell, 2011. 19(2): p. 244–56.

7. Sutherland, K.D., A.S. Ireland, and T.G. Oliver, Killing SCLC: insights into how to target a shapeshifting tumor. Genes Dev, 2022. 36(5-6): p. 241–258.

8. Zhang, W., et al., Small cell lung cancer tumors and preclinical models display heterogeneity of neuroendocrine phenotypes. Transl Lung Cancer Res, 2018. 7(1): p. 32–49.

9. George, J., et al., Comprehensive genomic profiles of small cell lung cancer. Nature, 2015. 524(7563): p. 47–53.

10. van Groningen, T., et al., Neuroblastoma is composed of two super-enhancer-associated differentiation states. Nat Genet, 2017. 49(8): p. 1261–1266.

11. van Groningen, T., et al., A NOTCH feed-forward loop drives reprogramming from adrenergic to mesenchymal state in neuroblastoma. Nat Commun, 2019. 10(1): p. 1530.

12. Boeva, V., et al., Heterogeneity of neuroblastoma cell identity defined by transcriptional circuitries. Nat Genet, 2017. 49(9): p. 1408–1413.

13. Balanis, N.G., et al., Pan-cancer Convergence to a Small-Cell Neuroendocrine Phenotype that Shares Susceptibilities with Hematological Malignancies. Cancer Cell, 2019. 36(1): p. 17–34 e7.

14. Pearson, J.D., et al., Binary pan-cancer classes with distinct vulnerabilities defined by pro-or anti-cancer YAP/TEAD activity. Cancer Cell, 2021. 39(8): p. 1115–1134 e12.

15. Cai, L., et al., Cell-autonomous immune gene expression is repressed in pulmonary neuroendocrine cells and small cell lung cancer. Commun Biol, 2021. 4(1): p. 314.

16. Jiang, L., et al., Genomic Landscape Survey Identifies SRSF1 as a Key Oncodriver in Small Cell Lung Cancer. PLoS Genet, 2016. 12(4): p. e1005895.

17. Breuer, K., et al., InnateDB: systems biology of innate immunity and beyond--recent updates and continuing curation. Nucleic Acids Res, 2013. 41(Database issue): p. D1228–33.

18. Gazdar, A.F., et al., Characterization of variant subclasses of cell lines derived from small cell lung cancer having distinctive biochemical, morphological, and growth properties. Cancer Res, 1985. 45(6): p. 2924–30.

19. Tumilowicz, J.J., et al., Definition of a continuous human cell line derived from neuroblastoma. Cancer Res, 1970. 30(8): p. 2110–8.

20. Canadas, I., et al., Targeting epithelial-to-mesenchymal transition with Met inhibitors reverts chemoresistance in small cell lung cancer. Clin Cancer Res, 2014. 20(4): p. 938–50.

21. Ayers, M., et al., IFN-gamma-related mRNA profile predicts clinical response to PD-1 blockade. J Clin Invest, 2017. 127(8): p. 2930–2940.

22. Montoro, D.T., et al., A revised airway epithelial hierarchy includes CFTR-expressing ionocytes. Nature, 2018. 560(7718): p. 319–324.

23. Plasschaert, L.W., et al., A single-cell atlas of the airway epithelium reveals the CFTR-rich pulmonary ionocyte. Nature, 2018. 560(7718): p. 377–381.

24. Chan, J.M., et al., Single cell profiling reveals novel tumor and myeloid subpopulations in small cell lung cancer. bioRxiv, 2020.

25. Davis, S. and P.S. Meltzer, GEOquery: a bridge between the Gene Expression Omnibus (GEO) and BioConductor. Bioinformatics, 2007. 23(14): p. 1846–7.

26. Ghandi, M., et al., Next-generation characterization of the Cancer Cell Line Encyclopedia. Nature, 2019. 569(7757): p. 503–508.

27. Pugh, T.J., et al., The genetic landscape of high-risk neuroblastoma. Nat Genet, 2013. 45(3): p. 279–84.

28. Vivian, J., et al., Toil enables reproducible, open source, big biomedical data analyses. Nat Biotechnol, 2017. 35(4): p. 314–316.

29. Rudin, C.M., et al., Comprehensive genomic analysis identifies SOX2 as a frequently amplified gene in small-cell lung cancer. Nat Genet, 2012. 44(10): p. 1111–6.

30. Subramanian, A., et al., Gene set enrichment analysis: a knowledge-based approach for interpreting genome-wide expression profiles. Proc Natl Acad Sci U S A, 2005. 102(43): p. 15545–50.

31. R Development Core Team, R: A language and environment for statistical computing, in Vienna, Austria. 2020, R Foundation for Statistical Computinng.

