## supplementary figures 1-9 for "The small cell lung cancer neuroendocrine transdifferentiation explorer"

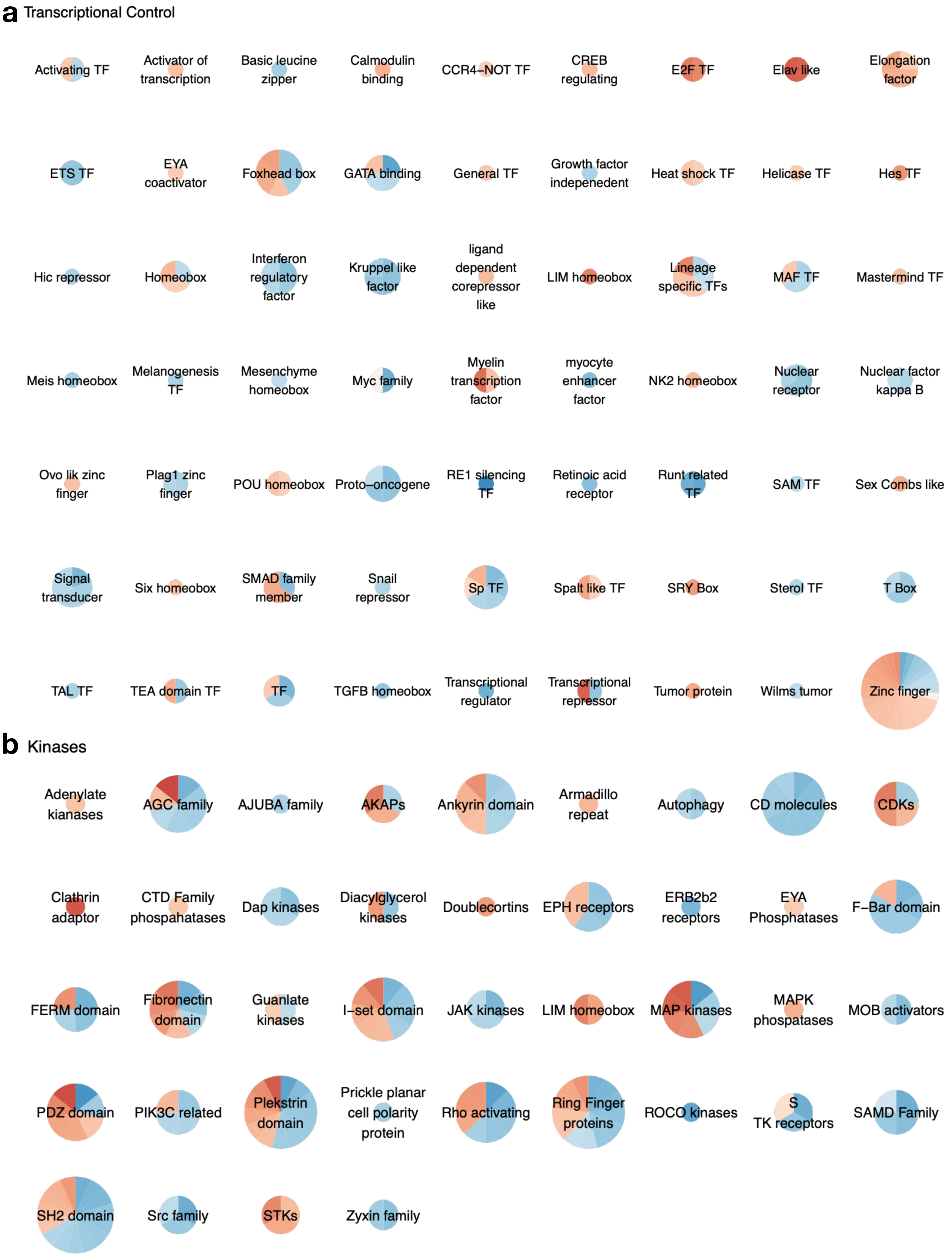
a Transcriptional Control
Activating TF
Activator of transcription
Basic leucine zipper
Calmodulin binding
CCR4–NOT TF
CREB regulating
E2F TF
Elav like
Elongation factor
ETS TF
EYA coactivator
Foxhead box
GATA binding
General TF
Growth factor indepenedent
Heat shock TF
Helicase TF
Hes TF
Hic repressor
Homeobox
Interferon regulatory factor
Kruppel like factor
ligand dependent corepressor like
LIM homeobox
Lineage specific TFs
MAF TF
Mastermind TF
Meis homeobox
Melanogenesis TF
Mesenchyme homeobox
Myc family
Myelin transcription factor
myocyte enhancer factor
NK2 homeobox
Nuclear receptor
Nuclear factor kappa B
Ovo lik zinc finger
Plag1 zinc finger
POU homeobox
Proto–oncogene
RE1 silencing TF
Retinoic acid receptor
Runt related TF
SAM TF
Sex Combs like
Signal transducer
Six homeobox
SMAD family member
Snail repressor
Sp TF
Spalt like TF
SRY Box
Sterol TF
T Box
TAL TF
TEA domain TF
TF
TGFB homeobox
Transcriptional regulator
Transcriptional repressor
Tumor protein
Wilms tumor
Zinc finger
b Kinases
Adenylate kianases
AGC family
AJUBA family
AKAPs
Ankyrin domain
Armadillo repeat
Autophagy
CD molecules
CDKs
Clathrin adaptor
CTD Family phospahatases
Dap kinases
Diacylglycerol kinases
Doublecortins
EPH receptors
ERB2b2 receptors
EYA Phosphatases
F–Bar domain
FERM domain
Fibronectin domain
Guanlate kinases
I–set domain
JAK kinases
LIM homeobox
MAP kinases
MAPK phospatases
MOB activators
PDZ domain
PIK3C related
Plekstrin domain
Prickle planar cell polarity protein
Rho activating
Ring Finger proteins
ROCO kinases
S TK receptors
SAMD Family
SH2 domain
Src family
STKs
Zyxin family

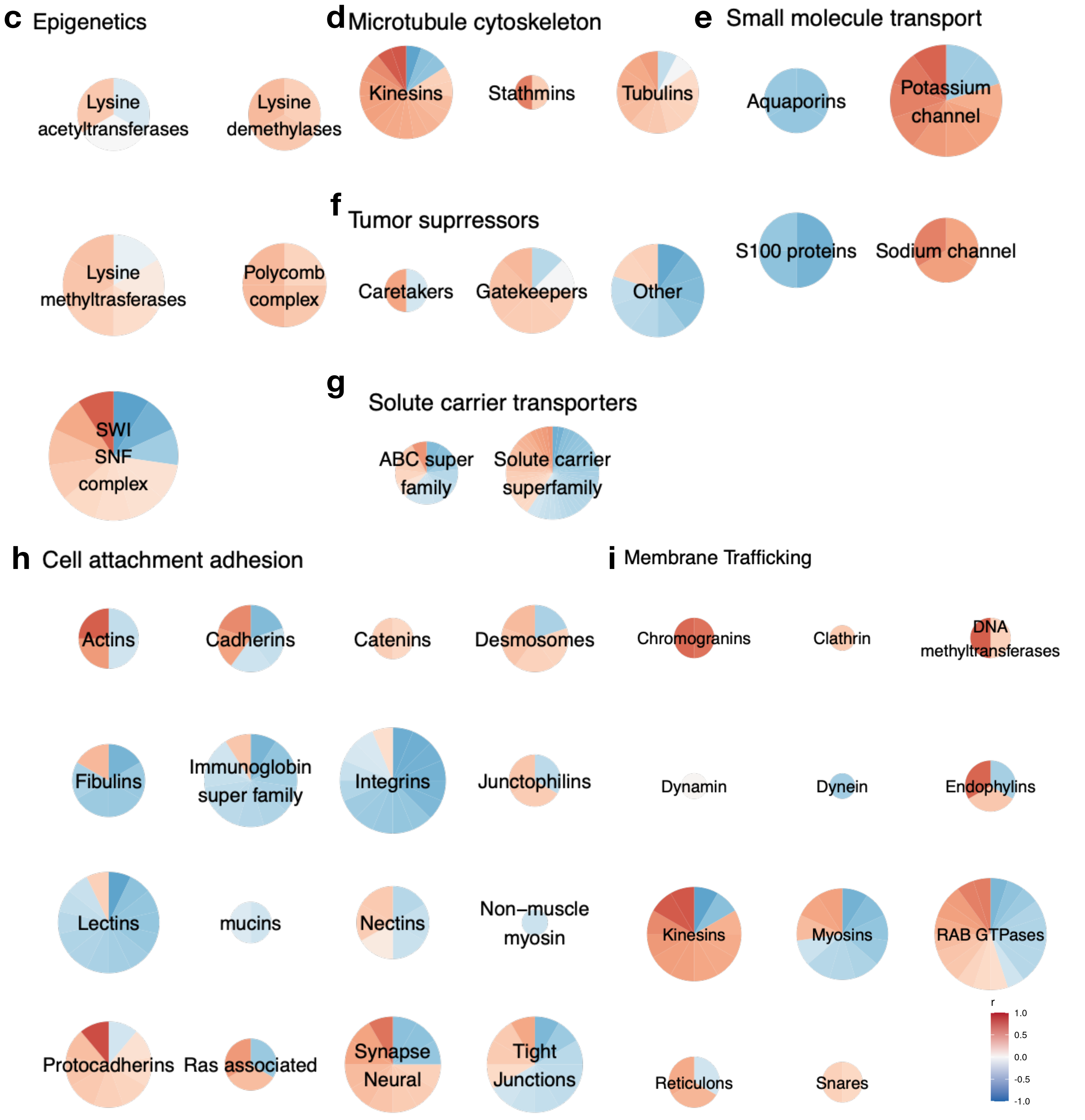


**Figure S1. Subgroups from nine gene families/pathways**
NE score vs. gene expression Pearson correlation profile is summarized for genes from different subgroups. Pie sizes are proportional to the number of genes within the set.

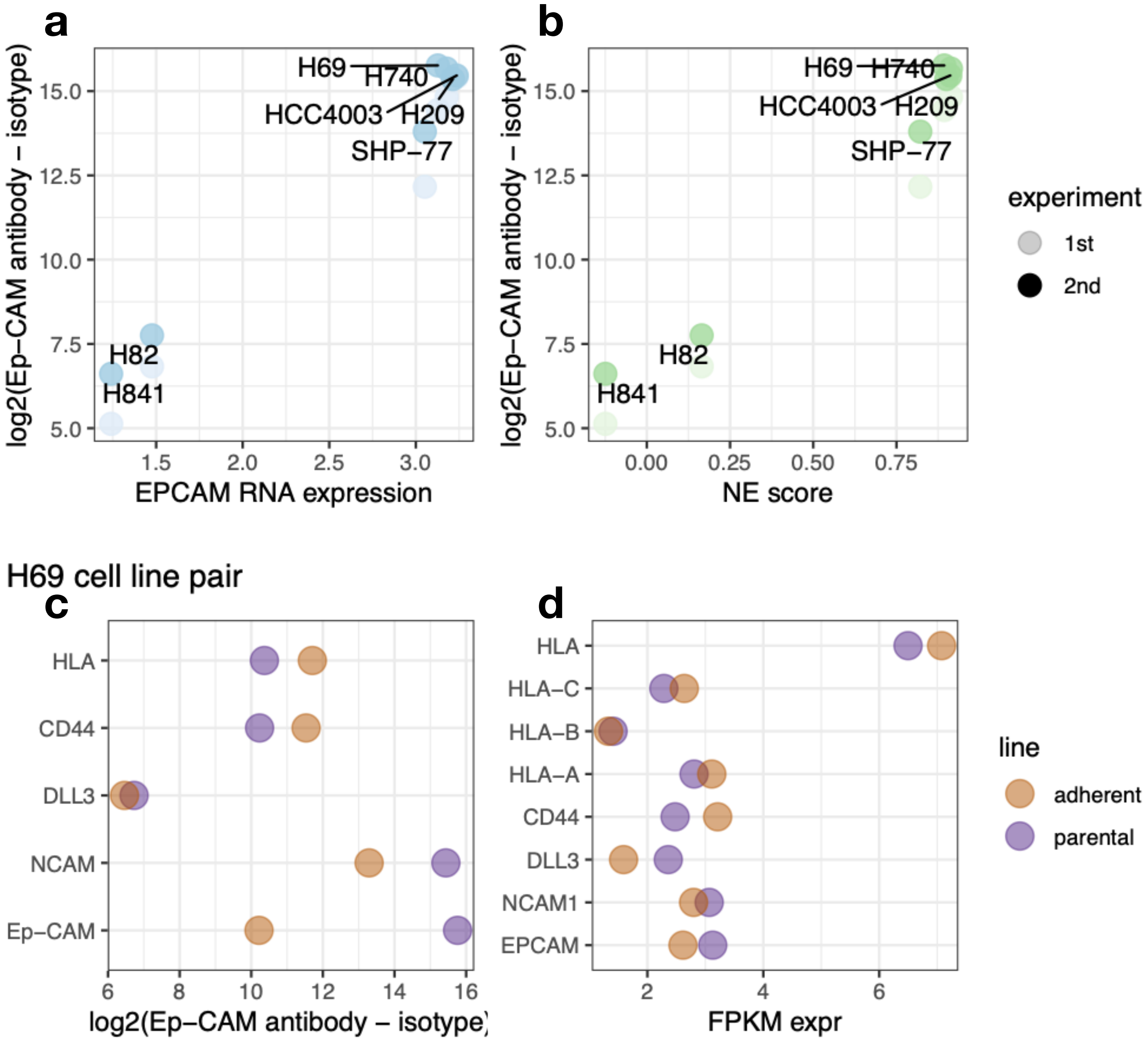


**Figure S2. NE and non-NE marker expression of SCLC cell lines**

**a.** Ep-CAM protein expression quantified by flow cytometry correlates well with Ep-CAM RNA expression. **b**. Ep-CAM expression is high in high-NE score SCLC cell lines and low in low-NE score cell lines. **c**. Decrease of NE marker expression (Ep-CAM, NCAM, and DLL3) and increase in non-NE marker expression (CD44 and HLA) in the adherent H69 subline compared to the parental H69 line measured by flow cytometry. **d**. As assessed by RNA-seq, decrease in NE markers (Ep-CAM, NCAM, and DLL3) and increase in non-NE markers (CD44, HLA-A, and HLA-C) in the adherent H69 subline compared to the parental H69 line.

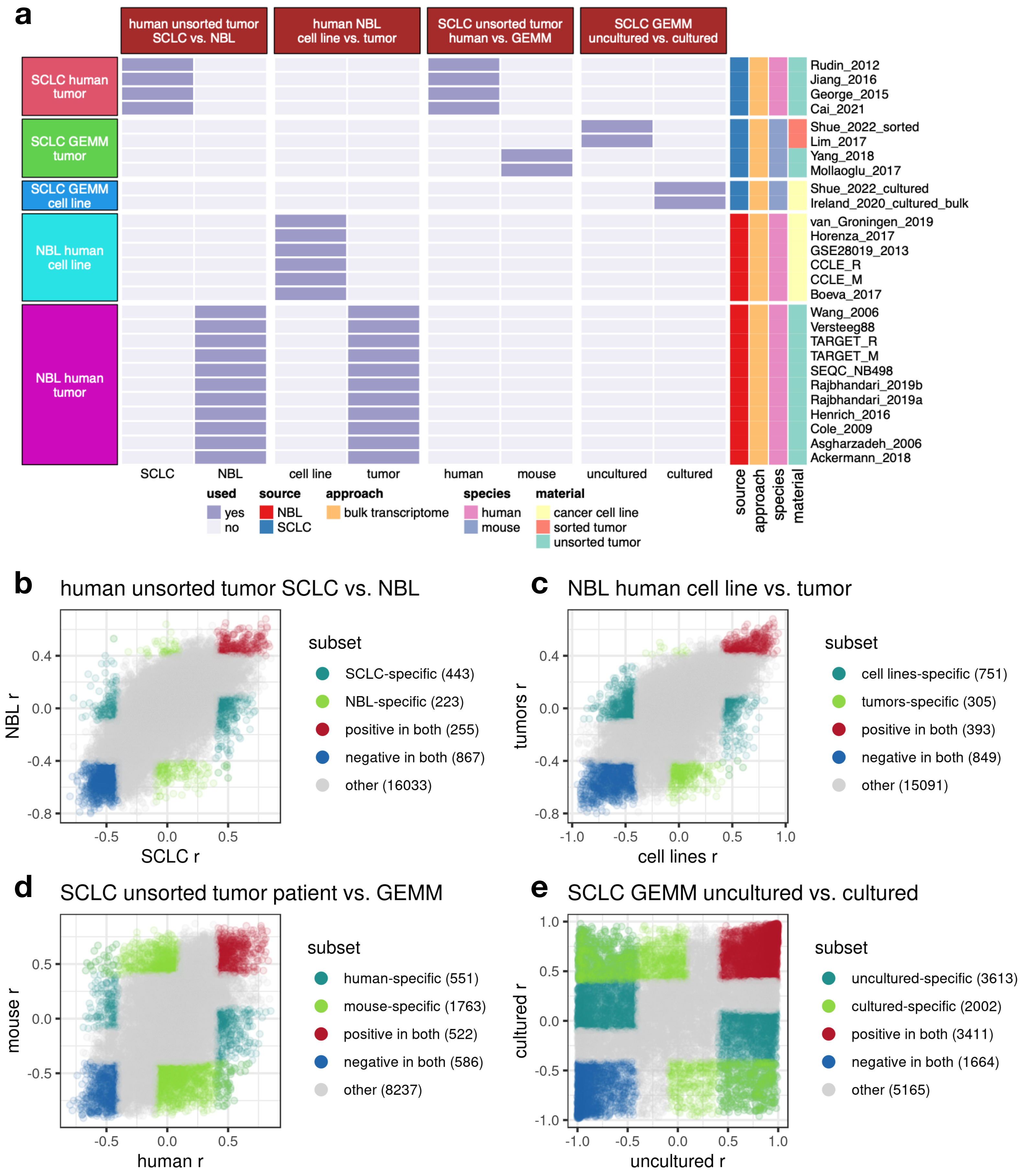


**Figure S3. Comparisons to dissect context-dependent NE associations**

**a.** Comparison scheme for examining context-specific NE associations. Four sets of comparisons in the top boxes. Datasets included in the meta-analysis are highlighted in the cells. Left-side box annotation follows the cluster assignment determined by Figure 2. Right-side color annotation denotes properties of each datasets. **b-e.** Scatterplots comparing NE score - gene expression correlations in different datasets. X- and y- axes values are Pearson correlation coefficient (r) summarized by meta-analysis. Context-specific genes are defined as those with r > .0.4 or < -0.4 in one meta-analysis and absolute r < 0.1 or with opposite direction in the other meta-analysis. The number of genes under different classes are indicated in brackets.

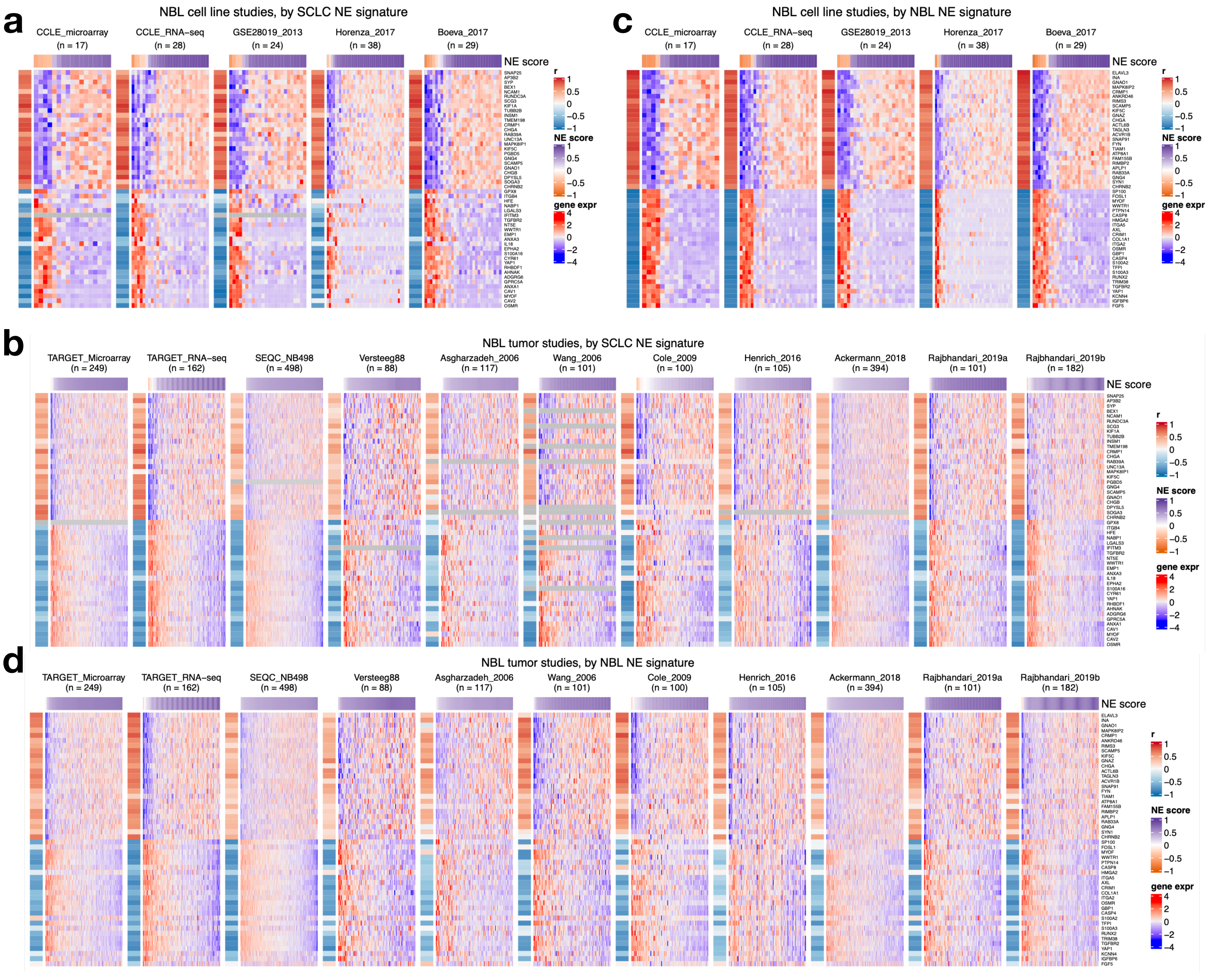


**Figure S4. An NBL-specific NE signature for NE score computation in NBL samples**

**a**-**b**, Heatmaps of SCLC NE gene signature in NBL cell lines (**a**) and tumor (**b**) datasets. **c**-**d**, Heatmaps of NBL NE gene signature in NBL cell lines (**c**) and tumor (**d**) datasets. In each heatmap, left-side column denotes Pearson correlation between NE score and gene expression for each gene, top annotation denotes NE score for each sample. NBL-specific NE signature used in **c** and **d** can be found in supplementary information 6.

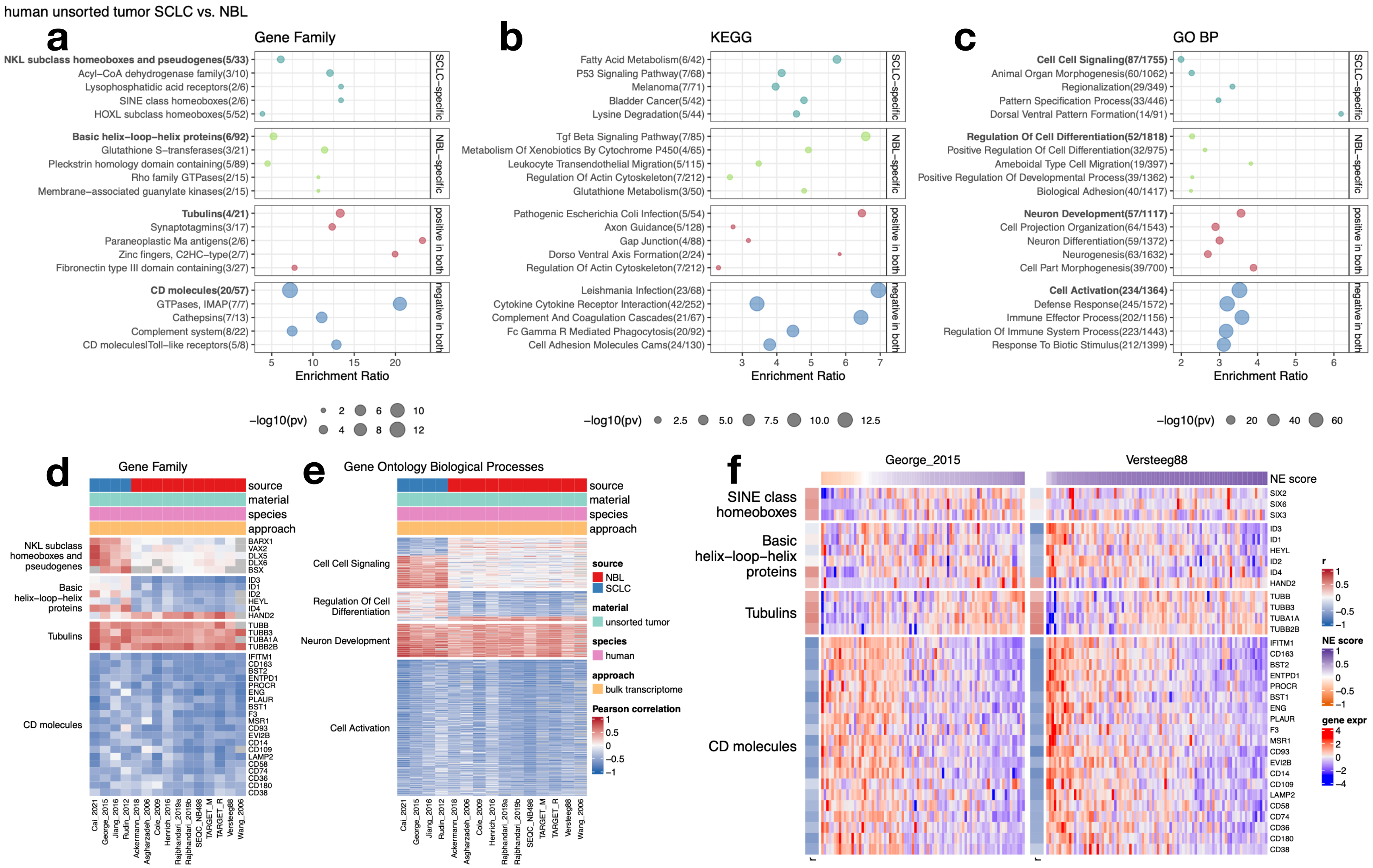


**Figure S5. Comparison between human SCLC and NBL tumors**

**a-c**. Geneset enrichment analysis results assessing pathways uniquely or commonly associated with NE scores in SCLC or NBL. Results for three gene set libraries - Gene Family (**a**), KEGG pathways (**b**), and Gene Ontology Biological Processes (GO BP) (**c**) are shown. More results and details can be found in **supplementary information 2**. Genesets in rectangles were further examined. **d-e**. Correlation heatmaps visualizing gene members from the top genesets from Gene Family (**d**), and GO BP (**e**) in four SCLC and eleven NBL human tumor studies. **f**. Gene expression heatmap visualizing relationship between NE scores and the expression of selected genes from top pathways in representative SCLC dataset (George_2015) and NBL dataset (Versteeg88) .

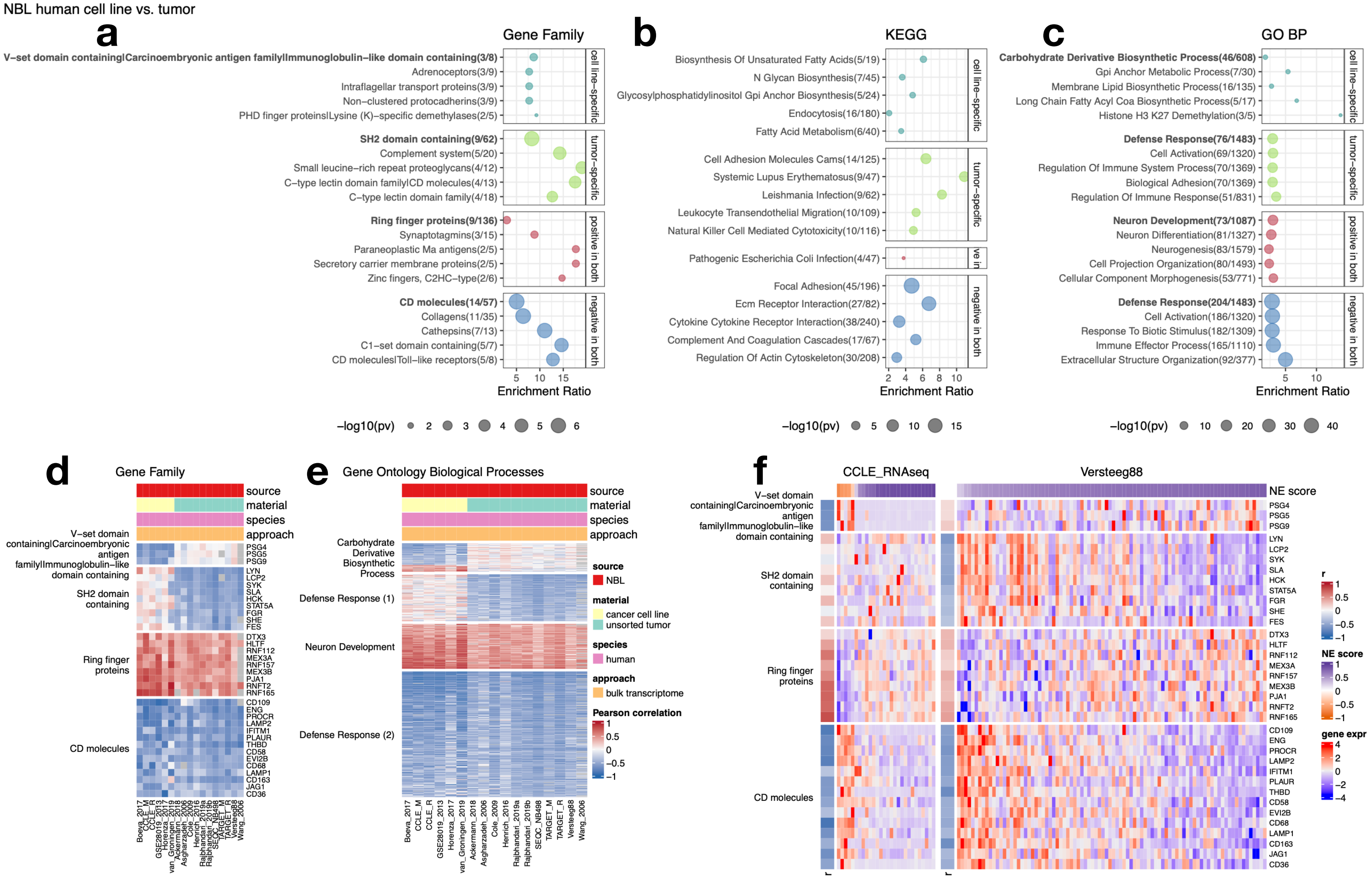


**Figure S6. Comparison between human NBL cell lines and tumors**

**a**-**c**. Pathway enrichment analysis results assessing pathways uniquely or commonly associated with NE scores in NBL cell lines or tumors. Results for three gene set libraries - Gene Family (**a**), KEGG pathways (**b**), and Gene Ontology Biological Processes (GO BP, **c**) are shown. More results and details can be found in **supplementary information 3**. Genesets in rectangles were further examined. **d**-**e**. Correlation heatmaps visualizing gene members from the top genesets from Gene Family (**d**), and GO BP (**e**) in six NBL cell and eleven NBL human tumor studies. **f**. Gene expression heatmap visualizing relationship between NE scores and the expression of selected genes from top pathways in representative NBL cell line dataset (CCLE_RNAseq) and NBL tumor dataset (Versteeg88) .

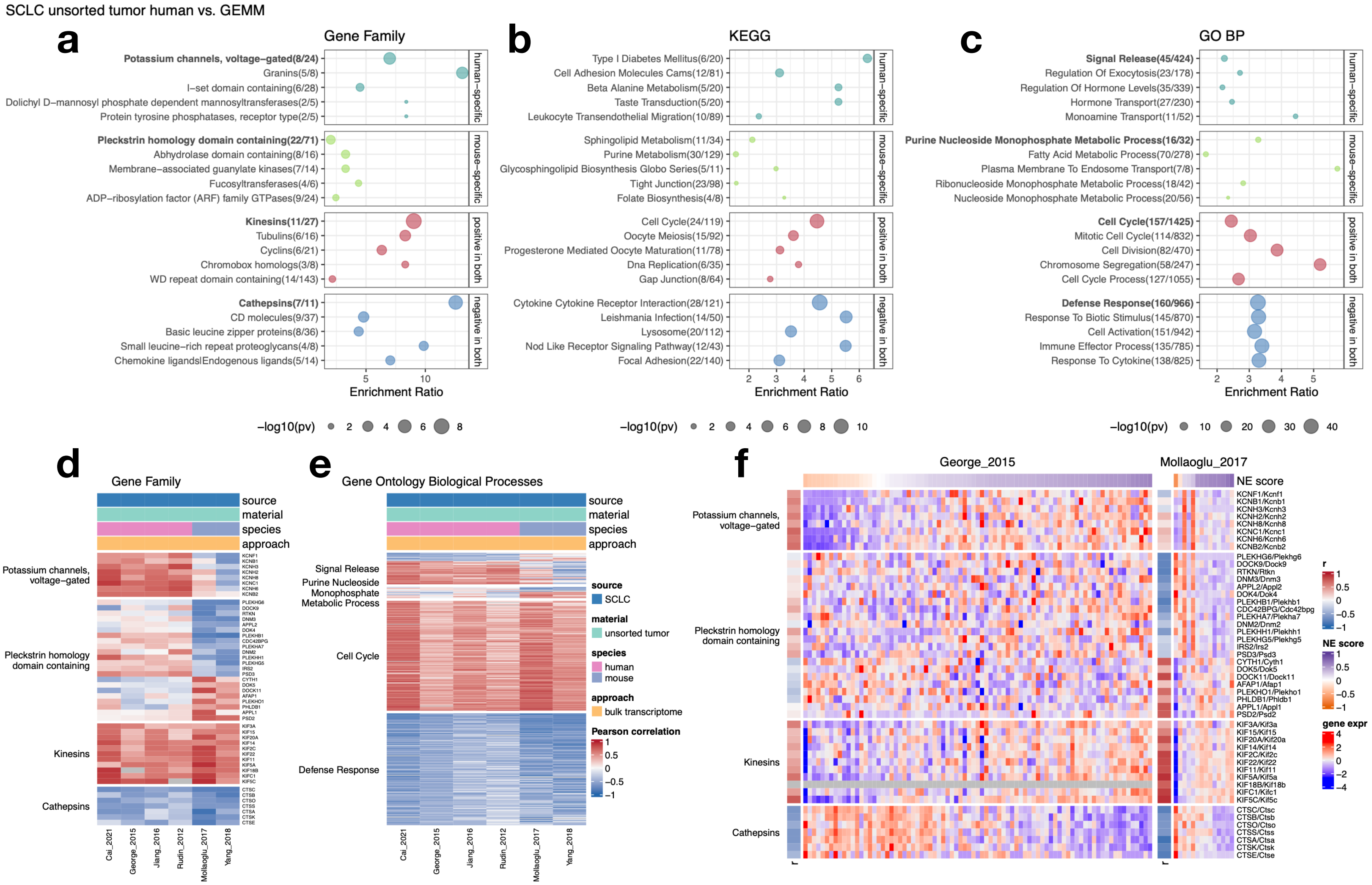


**Figure S7. Difference and similarity between SCLC human and GEMM tumors**

**a-c**. Pathway enrichment analysis results assessing pathways uniquely or commonly associated with NE scores in SCLC human tumors or GEMM tumors. Results for three gene set libraries - Gene Family (**a**), KEGG pathways (**b**), and Gene Ontology Biological Processes (GO BP) (**c**) are shown. More results and details can be found in **supplementary information 4**. Genesets in rectangles were further examined. **d-e**. Correlation heatmaps visualizing gene members from the top genesets from Gene Family (**d**), and GO BP (**e**) in four SCLC human tumor and three SCLC GEMM tumor studies. **f**. Gene expression heatmap visualizing relationship between NE scores and the expression of selected genes from top pathways in representative SCLC human tumor dataset (George_2015) and SCLC GEMM tumor dataset (Mollaoglu_2017) .

SCLC GEMM uncultured vs. cultured

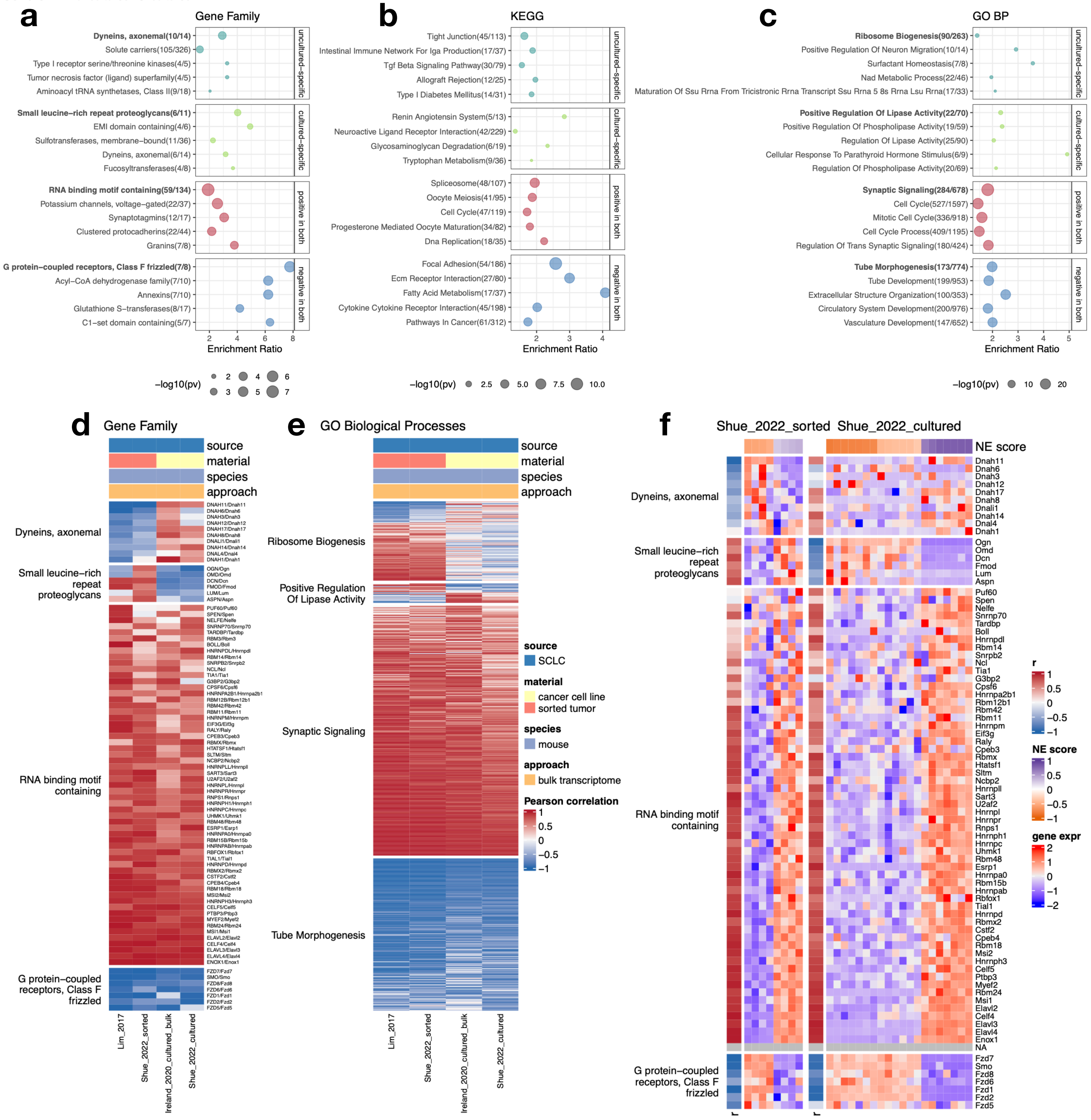


**Figure S8. Difference and similarity between SCLC GEMM uncultured and cultured cancer cells**

**a-c**. Pathway enrichment analysis results assessing pathways uniquely or commonly associated with NE scores in SCLC GEMM uncultured or cultured cancer cells. Results for three gene set libraries - Gene Family (**a**), KEGG pathways (**b**), and Gene Ontology Biological Processes (GO BP) (**c**) are shown. More results and details can be found in **supplementary information 5**. Genesets in rectangles were further examined. **d-e**. Correlation heatmaps visualizing gene members from the top genesets from Gene Family (**d**), and GO BP (**e**) in two SCLC GEMM uncultured cancer cell and three SCLC GEMM cultured cancer cell line studies. **f**. Gene expression heatmap visualizing relationship between NE scores and the expression of selected genes from top pathways in representative SCLC GEMM uncultured dataset (Shue_2022_sorted) and SCLC GEMM cultured dataset (Shue_2022_cultured).

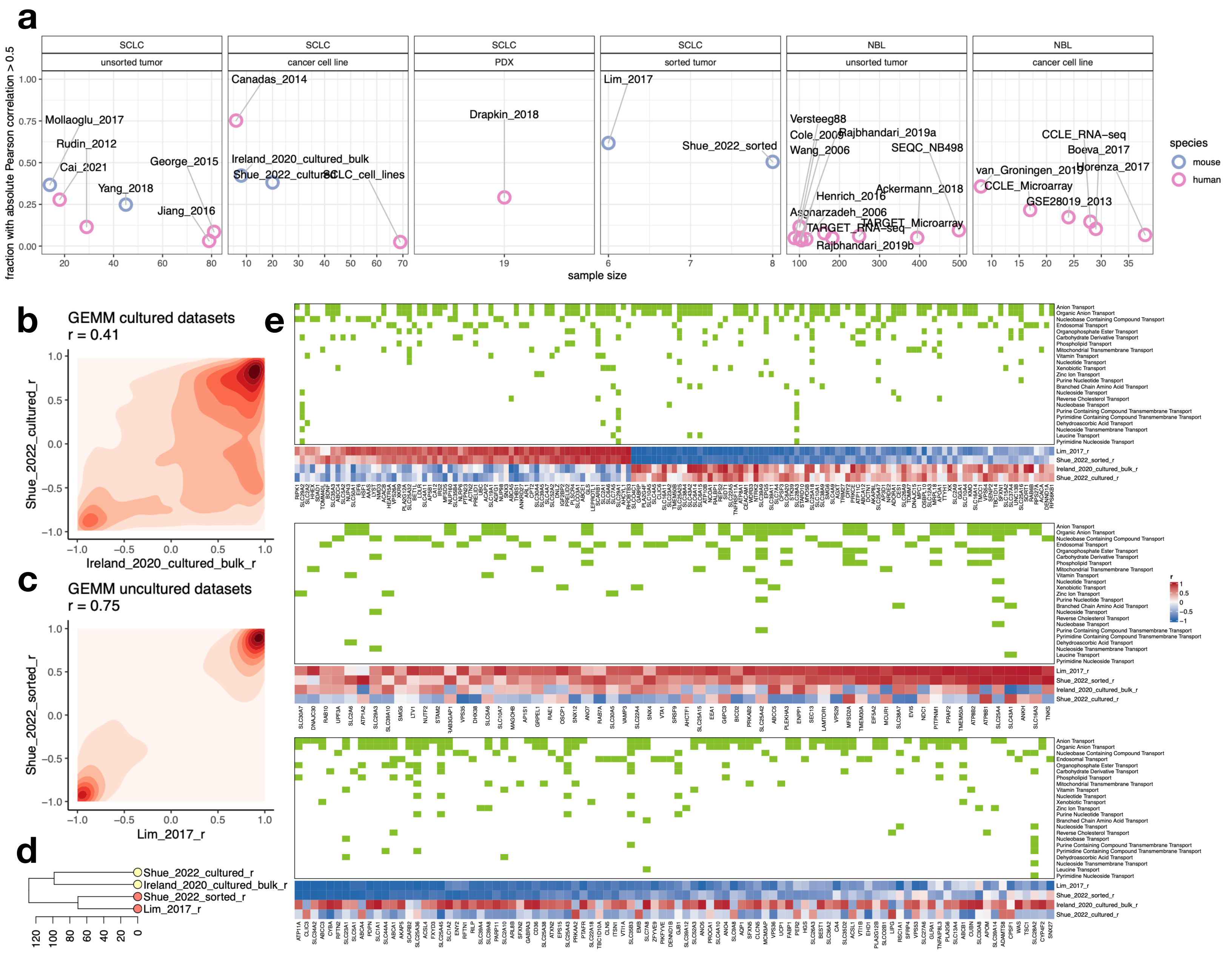


**Figure S9. Investigation of gene - NE score associations specific to culture conditions with SCLC GEMM datasets**
**a**. Fraction of genes with large effect size in their correlation with NE scores summarized for 30 datasets. NBL tumor datasets generally have fewer robustly correlated genes whereas studies with NE and non-NE groups have more robust correlations.
**b-c**. 2D density plot showing distribution of transcriptome NE score correlations in SCLC GEMM cultured datasets (**b**) and uncultured datasets (**c**). Correlation between the two cultured datasets are weaker than between the two uncultured datasets. **d**. Hierarchical clustering by gene - NE score association cluster SCLC GEMM datasets by culture status. **e**. Genes involved in metabolite transport that show uncultured/culture-specific NE score associations. Top: genes with better consistency between the two cultured datasets. Middle/bottom: genes with less consistency between cultured datasets. Geneset memberships are denoted on top of the heatmap.
